# Tautomerization constrains the accuracy of codon-anticodon decoding

**DOI:** 10.1101/2020.10.19.344408

**Authors:** Andriy Kazantsev, Zoya Ignatova

## Abstract

G○U(T) mismatch has the highest contribution to the error rate of base pair recognition in replication, as well as in codon-anticodon decoding in translation. Recently, this effect was unambiguously linked to keto-enol tautomerization, which enables the Watson-Crick (WC) geometry of the base pair. Structural studies of the ribosome revealing G○U in the WC geometry in the closed state of the A-site challenge the canonical induced-fit model of decoding and currently lack a physicochemical explanation.

Using computational and theoretical methods, we address effects of the ribosomal A-site on the wobble↔WC tautomerization reaction in G○U (wb-WC reaction), and the consequent implications for the decoding mechanism in translation. The free energy change of the wb-WC reaction in the middle codon-anticodon position was calculated with quantum-mechanical/molecular-mechanical umbrella sampling simulations. The wb-WC reaction was endoergic in the open A-site, but exoergic in the closed state. This effect can be explained in part by the decreased polarity of the closed A-site.

We developed a model of initial selection in translation that incorporates the wb-WC reaction parameters in the open and closed states of the A-site. In the new model, the exoergic wb-WC reaction is kinetically restricted by the decoding rates, which explains the observations of the WC geometry at equilibrium conditions. Moreover, the model reveals constraints imposed by the exoergic wb-WC reaction on the decoding accuracy: its equilibration counteracts the favorable contribution from equilibration of the open-closed transition. The similarity of the base-pair recognition mechanism in DNA polymerases allows extending this model to replication as well. Our model can be a step towards a general recognition model for flexible substrates.

## INTRODUCTION

The accuracy of base pair recognition during replication and translation has a fundamental role in cell fitness and evolution^1–4^. Some mismatches, such as G○U (G○T) mismatch, have particularly large contributions to error rate^5–10^. The tautomeric hypothesis by Watson and Crick^11,12^ suggests that a source of base pair recognition errors can be attributed to keto/enol and amino/imino tautomerism of nucleobases. G○U(T) base pair formed with the canonical keto states of the nucleobases is unable to adopt the Watson-Crick (WC) geometry and predominantly exits in the wobble (wb) geometry^13^. Formation of enol tautomeric states (G* or U*(T*)) enables the WC geometry, potentially leading to errors during base pair recognition, but the biologically-relevant mechanisms of this process remained elusive until recently.

Using computational chemistry methods, Brovarets’ and Hovorun predicted wobble↔WC tautomerization reaction in G○U(T) base pair^14,15^ (wb-WC reaction, Fig. 1A), which was experimentally confirmed by NMR in DNA and RNA duplexes in water solution^13^, where this reaction was endoergic (i.e. the wobble geometry was more populated than the WC)^13,16^. Recently, by incorporating tautomeric populations measured in DNA duplexes in water solution into a numerical kinetic model of DNA replication, a good agreement between predicted and experimentally observed misincorporation rates was obtained^16^. It might indicate a negligible role of the active site environment on the properties of the wb-WC reaction. However, both experimental^17^ and computational^18,19^ studies demonstrate stabilization of the WC geometry of G○T in the closed state of active site in several DNA polymerases, implicating a more complicated mechanism of base pair recognition. The WC geometry of G○U was also observed in the closed state of the decoding site (A-site) of the ribosome at the first two codon-anticodon positions: in crystal structures^20–23^ and later in Cryo-EM structures^24^.

**Figure 1:**
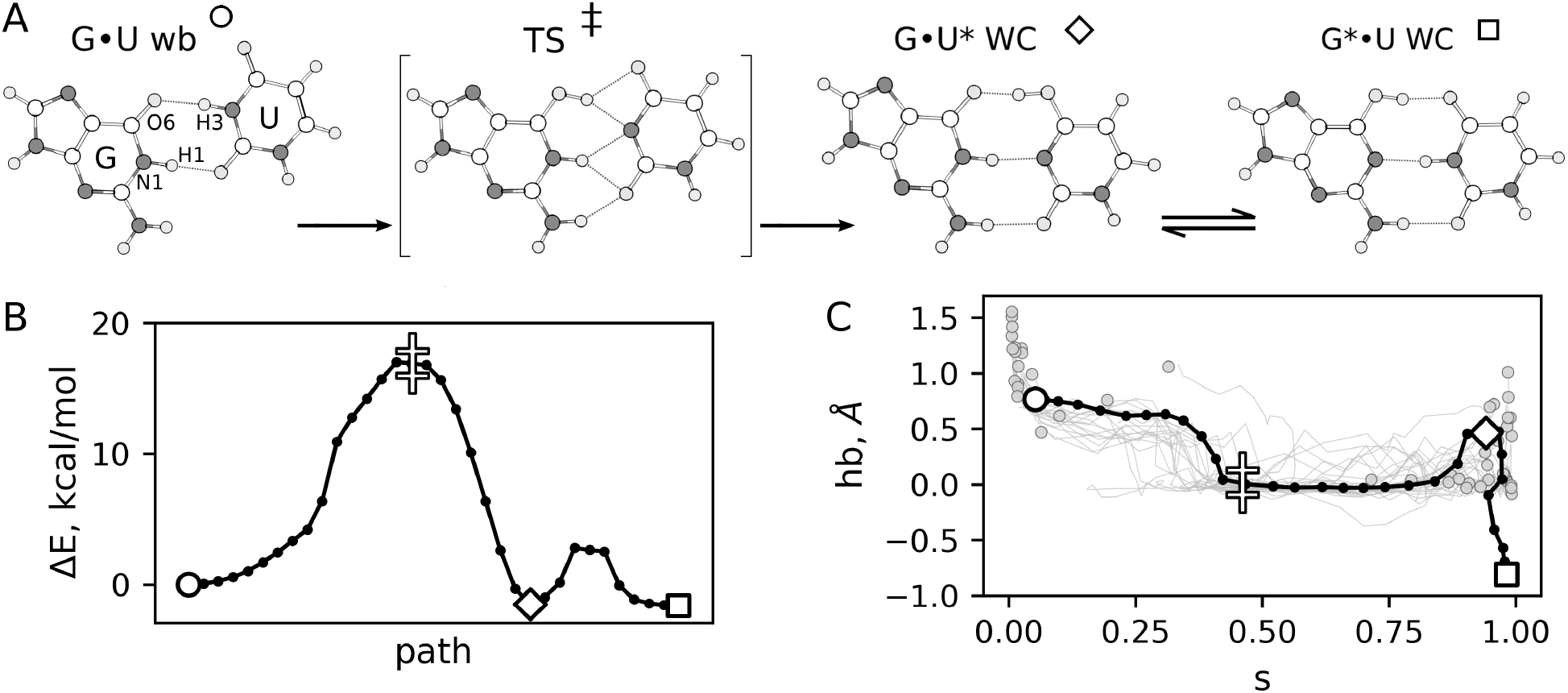
The wb-WC reaction. **A** – local minima and the transition state (TS) along the wb-WC reaction. Asterisk denotes enol tautomers. The markers shown at each state are used to denote the corresponding states in **B** – **C**. The wb-WC reaction consists of slow tautomerization reaction from G○U wb to G○U* WC, followed by fast double proton transfer reaction between G○U* and G*○U. **B** – reference potential energy profile of the wb-WC reaction calculated with nudge elastic band method (NEB) in gas phase. **C** – reference wb-WC reaction path from NEB, projected on the collective variables used in umbrella sampling calculations *s* (path collective variable) and *hb* (proton transfer variable). Gray lines and gray circles denote trajectories from the committor analysis, initiated from the TS

These observations of the WC geometry of G○U(T) challenge the classical model of substrate recognition in replication and translation - the induced-fit model, which assumes rigid substrates. The open↔closed conformational transition of the ribosomal decoding site, the molecular mechanism of the ‘‘induced fit”^24,25^, provides the highest decoding accuracy in the equilibrium limit, but is shifted from equilibrium due to its high forward rates, and the rates of the following irreversible reaction^26–29^. Theoretical studies based on the rigid substrate approximation have concluded that decoding in translation has been evolutionarily optimized for higher rate rather than for higher accuracy^27,28,30^. Since the rate constants affecting the open↔closed transition are not rate-limiting in translation elongation^27^, a possible advantage of such trade-off remains unclear and additionally challenges the induced-fit model. Hovorun and coworkers have suggested a non-equilibrium exoergic wb-WC reaction in the closed state of a DNA polymerase active site, kinetically restricted by the lifetime of the recognition state^15^. Such mechanism applied to the codon-anticodon decoding might solve the above described problems, but requires formalization and detailed analysis.

Clearly, studying environmental effects of the ribosomal decoding site on the wb-WC reaction is essential for understanding the decoding mechanism. In the absence of experimental methods suitable for addressing this problem, computational methods provide such opportunity. Until now, two computational studies have addressed nucleobase tautomerism in codon-anticodon decoding^31,32^, both of which argue against stabilization of the WC geometry in G○U at the decoding site, thus contradicting the experiments^20–24^. However, these computational studies used force fields for energy calculations, which may not be the most accurate approach to study tautomerization, especially when the rare tautomers were not parameterized, as in ref.^31^.

Here, we overcome the force-field-related biases by performing hybrid quantum-mechanical/ molecular-mechanical (QM/MM) simulations. We use QM/MM umbrella sampling (US) method to calculate free energy profiles of the wb-WC reaction in benchmark systems (DNA duplex in water, and a single base pair in benzene), ribosomal A-site models and active sites of DNA polymerases. Our calculations reveal positive free energy of the WC geometry formation *ΔG_wc_* in the model of the open A-site, and negative *ΔG_wc_* in the closed model, consistent with the structural studies^20–24^. We also found different effects of DNA polymerases on the wb-WC reaction: Δ*G_wc_* in the active site of polymerase *β* was negative, consistent with the structural studies^17^, while Δ*G_wc_* in DNA polymerase T7 was not significantly affected compared to the DNA duplex in water. We analyze dielectric constant of the studied environments to explain the observed effects on the wb-WC reaction.

To study implications of the exoergic wb-WC reaction in the closed A-site on the codon-anticodon decoding, we incorporated the wb-WC reaction into the model of initial selection in translation and derived analytical solutions for the new model. In this model, the exoergic wb-WC reaction is kinetically restricted by the residence time of the closed state of the decoding site. The new model is consistent with previously reported WC geometry of G○U in the structural studies at equilibrium conditions^20–24^. In addition to uniting structural and kinetic data, the model reveals constraints on decoding: the equilibration of the wb-WC reaction counteracts the equilibration of the open↔closed transition, thereby constraining the error rate of decoding.

The similarities with DNA polymerases suggest that the proposed model can be also applied to replication. The model derived here might be a step towards a more general model of recognition capable of describing flexible substrates.

## METHODS

### Details on the methods are provided in Supplementary Methods

#### QM calculations

We performed quantum chemistry (QM) calculations to obtain the reference dependence of the wb-WC equilibrium on the dielectric constant of a solvent, and to select the optimal QM level of theory for our QM/MM calculations. Using geometries of the wb-WC reaction minima and transition state (TS) optimized at density functional theory (DFT) levels *ω*B97X-D3/def2-TZVP, B3LYP-D3BJ/def2-TZVP and BLYP-D3BJ/def2-SVP in gas phase and implicit water model, we calculated single-point energies with multiple DFT and semiempirical (SE) methods. The methods were assessed based on the energies (*ΔE_wc_*) or enthalpies (*ΔH_wc_*) of the WC geometry formation (*ΔE_wc_* = *ΔE*(G*oU WC) - *ΔE*(G○U wb)). Our benchmark calculations showed PM7^33^ as an optimal QM method for our QM/MM approach, as it was able to accurately reproduce *ΔH_wc_* calculated with DFT (Table S2), while allowing vast sampling due to its low computational cost. The activation enthalpies obtained with PM7 were much less accurate (Table S2). Orca 4.2.1^34^ was used for all *ab initio* QM calculations, while MOPAC-2016^35^ was used for all SE calculations.

#### System setup and MD simulations

We studied the environmental effects on the wb-WC reaction in the following systems: DNA heptamer in water solution and a single G○T base pair in benzene (benchmark systems); ribosomal A-site models: closed, abasic and open; human DNA polymerase *β* (pol-*β*) and DNA polymerase from T7 virus (T7-pol).

DNA heptamer model was prepared from the solution NMR structure of DNA dodecamer (PDB ID: 1BJD), centered on one of the G○T wobble base pairs in the dodecamer. Benzene model was prepared by solvating a single G○T base pair from the DNA heptamer system in a benzene box.

pol-*β* model was prepared from the crystal structure of pol-*β* with a G○C base pair in the closed active site (PDB ID: 4KLF)^36^, in which we replaced dCTP with dTTP. T7-pol was prepared in a similar way using the crystal structure with PDB ID 1T7P ^37^ as the initial coordinates.

To prepare the A-site models, we used the crystal structure of *T.thermophilus* 70S ribosome bound to tRNA*^Thr^* in the A-site (PDB ID: 6GSK) with U○G base pair in the second codon (AUC) - anticodon (GGU) position^23^. All residues within 35 Å radius of the second codon nucleotide were selected for the model (Fig. 2A). The outer shell of the selected sphere was restrained in all simulations. In the “closed” (native) A-site model the rRNA residues A1492, A1493 and G540 were in *out* conformation, surrounding the codon-anticodon helix (Fig. 2B). In the “abasic” model the nucleobases of A1492, A1493 and G540 were deleted. In the “open” model, used only in QM/MM US simulations (see below), we used harmonic biases from Zeng et al.^32^ to restrain A1492 and A1493 to the *in* position. All Mg^2+^ ions from the ribosome crystal structure were deleted in the models.

**Figure 2:**
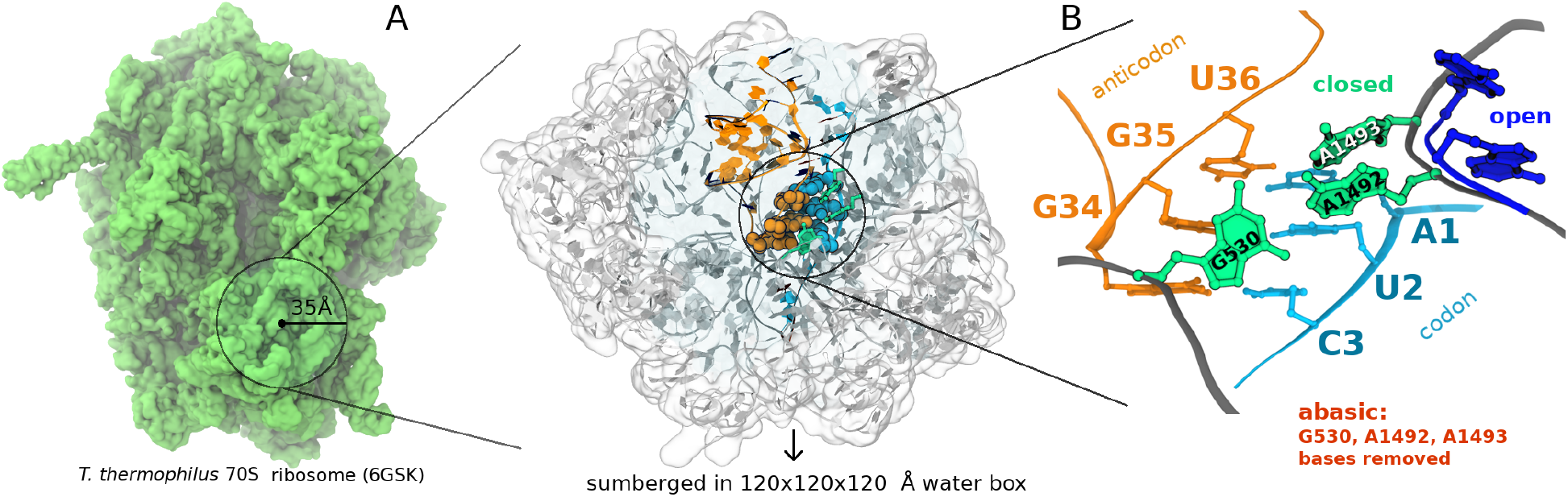
Preparation of the A-site models. **A** – visualization of the A-site model (right) and its approximate position and size in the 70S ribosome (left). **B** – close-up view on the codon-anticodon helix in the decoding center. The closed and the open A-site models differ by conformations of the rRNA residues A1492 and A1493 (green and blue in the closed and open states, respectively). In the abasic A-site model nucleobases of A1492, A1493 and G540 were deleted, leaving only the sugar-phosphate backbone.

Unless specified otherwise, all models were solvated in TIP3P^38^ water box with 0.15 M of NaCl. We employed a standard equilibration protocol (see Supplementary Methods), comprising steepest-descent minimization, solvent equilibration in NPT ensemble at standard conditions and gradual heating of the solutes to 298 K. Production MD simulations were performed in NVT ensemble using Langevin thermostat with 2 fs integration step. All MD simulations were performed using periodic boundary conditions. CHARMM36 force field^39–41^ in NAMD 2.12^42^ was used for all classical MD simulations.

#### Dielectric constant calculations

To analyze dielectric properties around the base pairs in the simulated models, we applied Kirkwood-Fröhlich formula (KFF), which relates dielectric constant (*ε*) to dipole moment fluctuations in a selected volume^43^. Using classical MD trajectories, we calculated *ε* of all nucleic acid, protein and water residues in spheres of radii ranging from 5 to 12 Å centered on the base pair of interest. Since *ε* calculated in a TIP3P water box using this approach depended on the probe volume radius until its convergence at ~ 20 ^Å^ (Fig. S3), the approach only allows a qualitative comparison between the studied systems.

#### QM/MM US simulations

We applied QM/MM umbrella sampling simulations (US) to calculate the free energy profiles (potential of mean force, PMF) of the wb-WC reaction in the models of ribosomal A-site and active site of DNA polymerases, as well as benchmark systems. All QM/MM calculations were performed on PM7/CHARMM36 level. The QM region comprised only nucleobases in a G○U(T) base pair of interest, with the QM-MM boundary placed at the glycosidic bonds. To select collective variables (CV) for the US calculations, we first optimized the wb-WC reaction path in G○U in gas phase using nudged elastic band (NEB) method at B3LYP-D3BJ/def2-TZVP level (Fig. 1B-C). The coordinate frames of this path became the reference for path collective variables^44^ (pathCV), which were used to calculate the position on the path (*s*) of the wb-WC reaction in the US simulations. *s* variable described the geometry change from wobble to WC, while a distance difference *hb* = d(O6-H3) - d(N1-H1) was used to describe proton transfers during the reaction. Selected CVs clearly distinguished all three local minima of the wb-WC reaction (Fig. 1C). Committor analysis was performed to verify the TS (Fig. 1C). Using *s* and *hb* as CVs, and adding a boundary potential to prevent large deviations from path *z*, we collected 100-400 ps of QM/MM US trajectories for each of 48 US windows per each system (~120 ns in total) and calculated PMF using weighted histogram analysis method (WHAM). We focused only on the relative energies and positions of local minima in the wb-WC reaction in our US simulations

#### Kinetic modeling

Decoding rate constants for a codon-anticodon combination with a G○U mismatch in the first position were obtained earlier by Pape et al.^45^, but only in the low-fidelity buffer conditions, in which the accuracy of the initial selection was extremely low (1.1), contrasting the *in vivo* measurements^5–9^. Therefore, for our kinetic modeling we selected decoding rate constants measured at high-fidelity conditions, which result in more realistic accuracy^46,47^. In the absence of high-fidelity measurements corresponding to a G○U mismatch, we used the set of rate constants corresponding to the A○C mismatch in the first codon-anticodon position^46,47^ Table S3. Rate constants 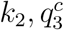 and 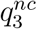 were assigned arbitrary. Numerical solutions of the kinetic model were obtained by numerical integration of ordinary differential equations (ODE) in Python. We describe derivations of analytical solutions in Appendix.

## RESULTS AND DISCUSSION

### Dielectric constant of base pair environments

It was speculated previously that closing of the ribosomal A-site, via conformational change of the rRNA residues A1492, A1493 and G540, desolvates the codon-anticodon helix, increasing base-pairing selectivity by an energy penalty from the lost H-bonds with water in mismatches^25,48^. We hypothesized that desolvation may also bring an opposite contribution to the accuracy of G○U mismatch recognition. Our benchmark calculations (Fig. S1) demonstrate that the wb-WC reaction is exoergic in gas phase and very nonpolar implicit solvents, corroborating previous computational studies^14,15,18,49^. We reasoned that a potentially decreased polarity of the closed ribosomal A-site may be responsible for the stabilization of the WC geometry of G○U observed in the structural studies.

We applied Kirkwood-Fröhlich formula (KFF) to qualitatively compare dielectric constant e of the decoding site environments between the closed and abasic models. In the latter model, nucleobases of residues A1492, A1493 and G540 were deleted (Fig. 2B). We observed a consistent *ε* decrease in the closed model for all codon-anticodon positions (Fig. 3). However, only at some distance cutoffs and positions the difference was statistically significant, which can be explained by the limited length and replica numbers of the MD trajectories (Fig. S4). From our KFF calculations we conclude that the A-site closing decreases *ε* of environment of all three codon-anticodon positions.

**Figure 3:**
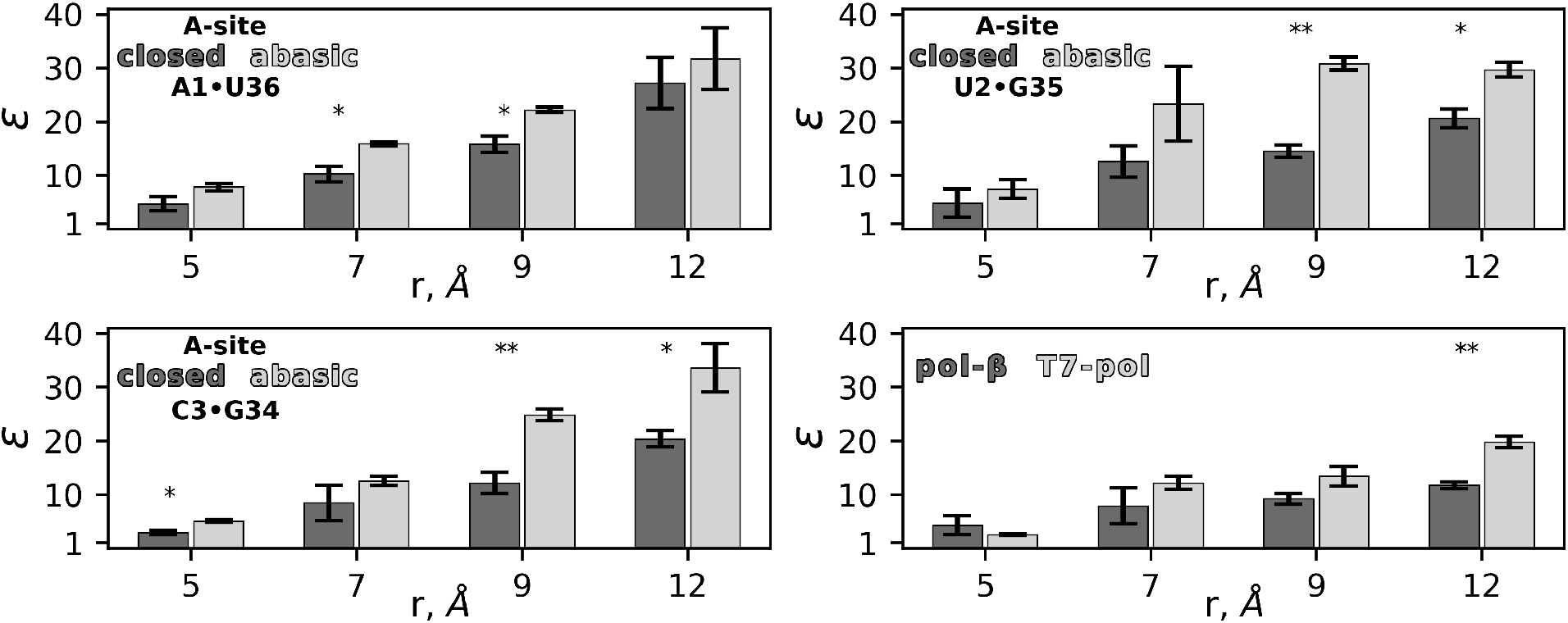
Dielectric constant *ε* of the base pair surroundings. **A**–**C** – *ε* difference between closed and abasic A-site models around each codon-anticodon base pair (A1-U36, U2-G35 and C3-G34) measured in spheres of radius r. **D** – *ε* difference between pol-*β* and T7-pol around the G○T base pair in the active site. Data are means *±* SD (n≥2 replica per model). *, *P* < 0.05; **, *P* < 0.01 (Student’s t-test)

We also compared the active sites of pol-*β* and T7-pol using the same approach. The active site environment in pol-*β* was generally less polar compared to T7-pol, but the difference was statistically significant only at 12 Å cutoff (Fig. 3).

In sum, our KFF calculations revealed decreased polarity in the closed compared to the abasic ribosomal A-site model, and in pol-*β* compared to T7-pol.

### Free energy change of the wb-WC reaction from QM/MM US

The effects we observed might indicate a putative environmental contribution on the energetics of the wb-WC reaction, but we cannot estimate the extent of this contribution on the wb-WC equilibrium. Therefore, we calculated potential of mean force (PMF) of the wb-WC reaction in the studied molecular environments using QM/MM umbrella sampling (US) calculations. The PMF from the converged part of the US trajectories of the DNA model is shown on Fig. 4A and the converged PMFs from the rest of the systems are shown on Fig. S5. The only quantitative property we derived from the PMFs was the total free energy change of the wb-WC reaction Δ*G_wc_* = Δ*G*(G*○U WC) - Δ*G*(G○U wb). Δ*G_wc_* was used to evaluate the convergence of the US simulations by calculating PMF separately for consecutive batches of ~ 20 ps.

**Figure 4:**
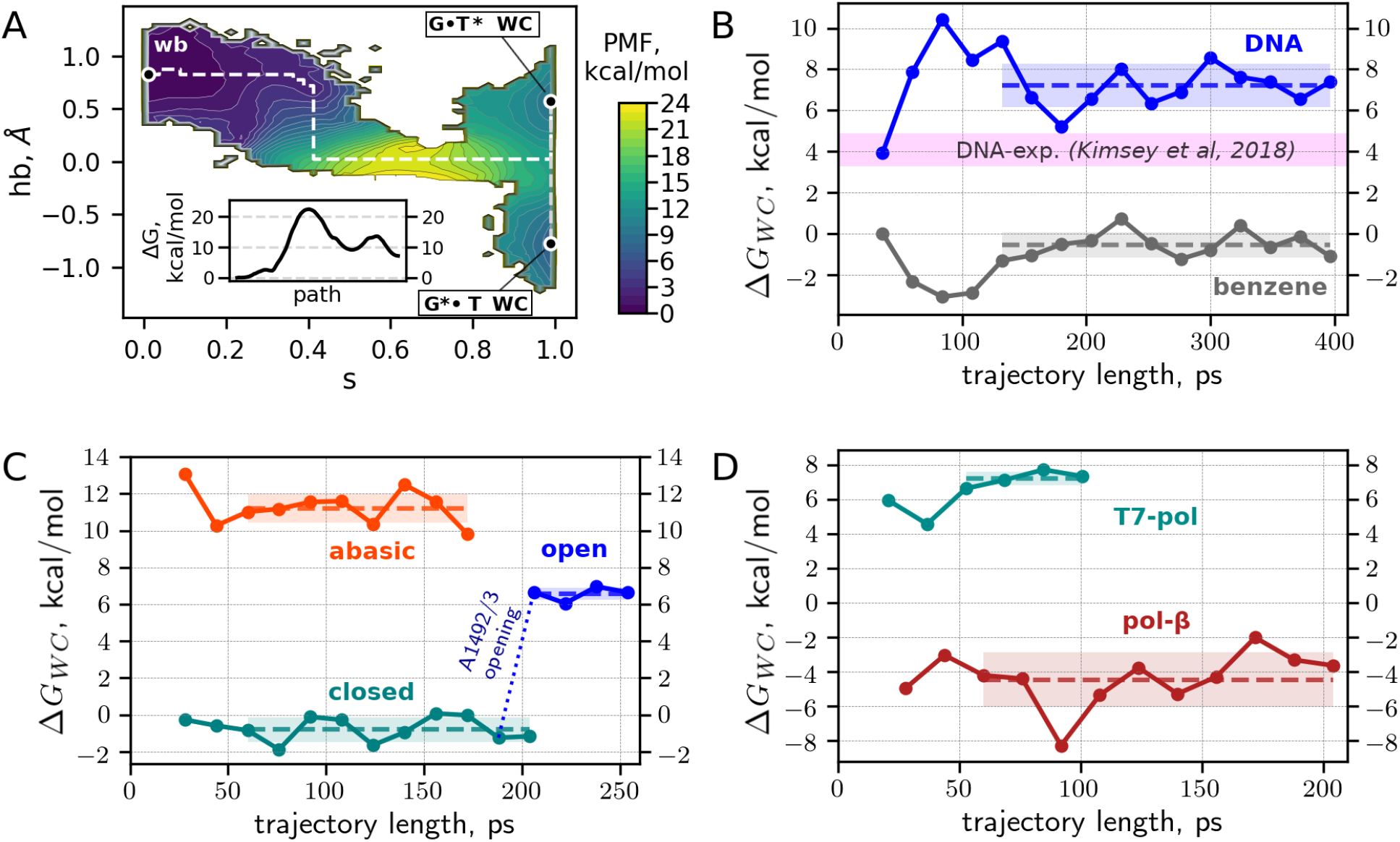
PMF from US calculations and its convergence. **A** – US-derived PMF of the wb-WC reaction in the DNA heptamer system. Black circles denote the minima corresponding to wb and WC structures. White dashed line denotes the minimal free energy path of the wb-WC reaction, from G○T wb to G*○T WC. The inset shows the free energy profile along the minimal free energy path. PMFs from the other six systems are shown on the Fig. S5. **B**–**D** – convergence of the Δ*G_wc_* in benchmark systems (**B**), A-site models with G○U in the middle codon position (**C**) and DNA polymerases (**D**). For each Δ*G_wc_* trajectory, the shaded area and the dashed line of the corresponding color denote the standard deviation and mean of the set of batches selected for convergence evaluation. Magenta-shaded area in **B** denotes the range of Δ*G_wc_* in DNA duplexes in solution measured by NMR^16^.

#### Benchmark systems

In order to verify the validity of our approach, we first applied it to benchmark systems: G○T base pair in the DNA heptamer duplex in water and a single G○T base pair in benzene. Δ*G_wc_* trajectories of the benchmark systems and the converged values of Δ*G_wc_* are shown in Fig. 4B and Table 1, respectively. After initial fluctuations from 4 to 10 kcal/mol, Δ*G_wc_* of the DNA model converged to 7.2 ± 1.1 kcal/mol. Our calculations somewhat overestimate the experimental Δ*G_wc_* range of 3.3-4.9 kcal/mol^16^, but are consistent with the recent computational result of ~ 6 kcal/mol derived from (*ω*B97X-D3/6-311G**)/AMBER calculations^18^. To assess the ability of our QM/MM setup to reproduce Δ*G_wc_* dependence on *ε* observed in implicit solvent models, we applied it to a single G○T base pair in explicit benzene environment (*ε* = 2.3). Δ*G_wc_* in benzene converged to −0.5 ± 0.6 kcal/mol, which is consistent with the QM calculations in implicit solvent model of similar *ε* (Fig. S1). In both benchmark systems the positions of the minima on the PMF were not qualitatively altered compared to the reference reaction path (Fig. S5, Fig. 1C). From our benchmark calculations we conclude that our setup is able to faithfully estimate environmental effects on the wb-WC reaction, reaching qualitative agreement with experiments and higher levels of QM theory.

**Table 1:** Δ*G_wc_* values from the US simulations

| System | $\Delta G_{wc}^a$ |
| --- | --- |
| DNA | $7.2 \pm 1.1$ |
| benzene | $-0.5 \pm 0.6$ |
| A-site closed | $-0.7 \pm 0.6$ |
| A-site abasic | $11.2 \pm 0.9$ |
| A-site open | $6.6 \pm 0.3$ |
| pol-β | $-4.4 \pm 1.6$ |
| T7-pol | $7.2 \pm 0.4$ |

<sup>a</sup> Mean $\Delta G_{wc} \pm$ standard deviation from the regions selected for convergence estimation on Fig. 4B-D, kcal/mol

#### Ribosomal A-site

We started the QM/MM US simulations of the A-site effects on the wb-WC reaction in the second codon-anticodon U○G base pair in the models of closed and abasic A-site. Δ*G_wc_* in the closed A-site converged to −0.7 ± 0.6 kcal/mol, while in the abasic A-site it converged to 11.2 ± 0.9 kcal/mol (Fig. 4C, Table 1). However, the abasic model is only a rough approximation of the open state of the A-site. To estimate effects of the rRNA nucleotides A1492 and A1493 more accurately, we created the “open” A-site model. In this model, harmonic restraints used in the previous studies^32^ were applied to change A1492 and A1493 conformations from *out* to *in* (Fig. 2B). G530 remained in the closed-like conformation in the open model. These restraints were applied to the coordinates from the end part of the closed state US trajectories and maintained for approximately 70 ps in each US window. Δ*G_wc_* in the open A-site converged to 6.6 ± 0.3 kcal/mol (Fig. 4C, Table 1). While in the abasic model the position of the wb minimum was not qualitatively altered compared to the reference reaction path (*s* = 0), in the closed and open models it shifted to higher values of the path variable (*s* ≈ 0.1), suggesting a slight destabilization effect on the wobble geometry, likely exerted by G530 (Fig. S5). This effect was not enough to shift the wb-WC equilibrium towards WC in the open model, indicating a critical role of A1492 and A1493 residues.

#### DNA polymerases

The previous QM/MM study have addressed effects of the DNA polymerase *λ* environment on the wb-WC reaction^18^. Here, we analyzed effects of two other DNA polymerases: low fidelity human DNA polymerase *β* (pol-*β*) and high-fidelity DNA polymerase from T7 virus (T7-pol). In both models, the wb-WC reaction was simulated in G○T base pair in the closed state of a polymerase active site. Δ*G_wc_* in pol-*β* converged to −4.4 ± 1.6 kcal/mol, indicating largely exoergic wb-WC reaction. This result corroborates previous structural studies, which reveal WC-like G○T base pair in the closed active site of pol-*β*^17^. Δ*G_wc_* in T7-pol converged to 7.2 ± 0.4 kcal/mol, resembling Δ*G_wc_* value in the DNA duplex (Fig. 4B,D, Table 1). To the best of our knowledge, WC-like G○T base pairs were never observed in the active site of T7-pol. Comparing PMFs from pol-*β* and T7-pol revealed a striking shift in the wb minimum position towards the higher value (*s* ≈ 0.2) of the path variable in pol-*β* (Fig. S5). We attribute this shift to steric effects in pol-*β* that constrain a base pair geometry in the closed active site.

In sum, using QM/MM US calculations, we demonstrated the exoergic wb-WC reaction in G○U at the middle codon-anticodon position in the closed A-site model, and the endoergic reaction in the open and abasic models. These results corroborate structural studies of the ribosome^20–24^. Decreased *ε* in the closed state of the decoding site can be one of the contributions to this effect. Shifts in the wb minimum position on the PMF in the closed and open models might indicate steric constraints on the base pair geometry. A similar effect was observed in pol-*β* active site, in which the wb-WC reaction was highly exoergic, in contrast to T7-pol, where the reaction equilibrium was not altered compared to the DNA duplex. More detailed studies are needed to delineate the environmental contributions on the properties of the wb-WC reaction.

### The wb-WC reaction in the kinetic model of decoding in translation

The exoergic wb-WC reaction in the closed A-site requires reconsideration of the current model of decoding in translation, as the previously suggested “energy penalty” from tautomerization^20–23^ cannot be the basis of selectivity. With the exoergic wb-WC reaction in equilibrium settings the error rate of G○U mismatches in the second codon position would be close to 1, but the error rate is 10^-2^ to 10^-3^ instead^5–9^. To reconcile the exoergic wb-WC reaction with the model of decoding, the wb-WC reaction should be out of equilibrium in the closed A-site. To study the contribution of this reaction in decoding, we incorporated it into the model of initial selection.

Starting from the classical kinetic model of initial selection (Fig. S6), explained in details in^29^, we introduced the wb-WC reaction by separating each near-cognate (nc) state into wb and WC states, leaving the cognate (c) branch unchanged (Fig. 5-A). Rate constants between nc-wb and nc-WC states correspond to the rate constants of the wb-WC reaction in a given environment of the A-site. To simplify the model for the sake of deriving analytical solutions, we assumed instant wb-shifted wb-WC equilibrium in states *C*2 (initial binding) and *C*3 (codon recognition in the open A-site). This approximation is well justified if, as suggested by our US calculations, the open state of the decoding site does not shift the equilibrium towards WC, and thus the equilibration time is negligible. In this approximation the wb-WC equilibrium in *C*2 does not affect the model and needed only for model completeness. The wb-WC reaction in C4 (codon recognition in the closed A-site) was modeled explicitly via its forward and reverse rate constants *k_f_, k_r_*. We used decoding rate constants measured previously at 20 °C^46,47^. The nc decoding rate constants in this set correspond not to a G○U mismatch, but to the A○C mismatch in the first position. This and other approximations preclude quantitative predictions of the error rate from the model. We analyzed effects of the wb-WC reaction on decoding that are not restricted to any particular combination of the decoding rate constants.

**Figure 5:**
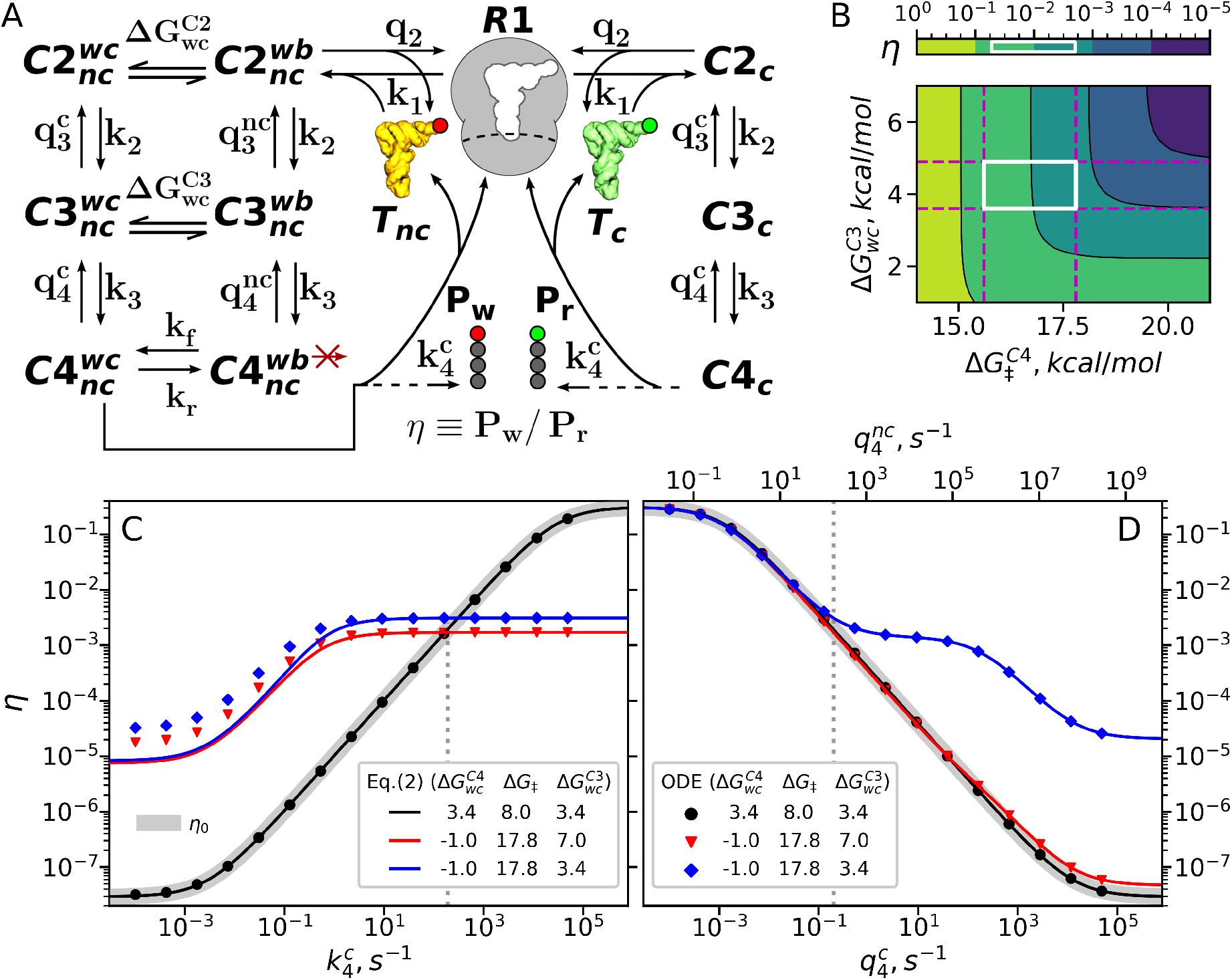
The wb-WC reaction in the kinetic model of decoding in translation. **A** – a scheme of the kinetic model of the initial selection in decoding with incorporated wb-WC reaction. Fig. S6 includes a detailed explanation of the decoding states and transitions. **B** – *η* as a function of Δ*G*_‡_ and 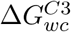, calculated using Eq. (2) for 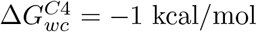. The highlighted region corresponds to the experimental range of Δ*G*_‡_ and Δ*G_wc_* in RNA duplexes^16^. **C-D** error rate as a function of 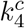 (**C**) and 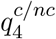 (**D**), calculated by Eq. (A.1) for the classical system (*η*_0_), and, for the specified wb-WC parameters, from numerical simulations (ODE) and by Eq. (2). 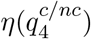 and 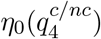 were obtained by varying 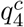 and 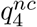 simultaneously at constant 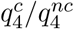 ratio. 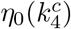 was also obtained at constant 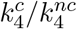 ratio. Δ*G* units in the plot legend are kcal/mol. The gray dotted vertical lines denote experimental values of 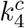 and 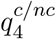. The deviation of Eq. (2)-predicted *η* from *η*(ODE) for low values of 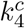 in **C** is due to increased *C*3 state population when the wb-WC reaction approaches equilibrium.

Inspired by the previously applied numerical kinetic model of selection in replication by Kimsey et al.^16^, we made the following assumptions: (i) nc-WC states are characterised with the same decoding rate constants as the cognate states, since the WC geometries are assumed to be indistinguishable for the decoding site; (ii) GTPase activation rate constant (*k*_4_) in 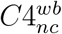 state is set to zero, assuming that the major contribution to this reaction rate comes from the nc-WC state, thus the contribution from nc-wb can be ignored; (iii) escape rate constants of the nc-wb states are taken from the classical nc states 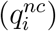 assuming that the cumulative escape rate is dominated by the nc-wb states, thus the nc→nc-wb rescaling can be neglected. Given the assumption (ii), the apparent near-cognate *k*_4_:

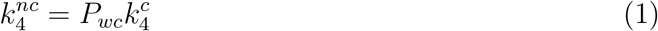

where *P_wc_* is the population of the 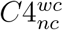 state - a function of the equilibrium WC populations in *C*3 and *C*4 states, and constraints imposed by the decoding rates on the reaction kinetics in *C*4. Using the equation for product concentration in a first-order reversible reaction, we derived (see Appendix) the equation for the error rate induced by the wb-WC reaction:

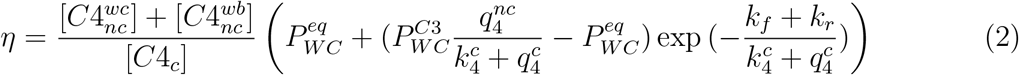

where 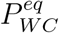 is the equilibrium WC population in *C*4 for a given (*k_f_, k_r_*), and 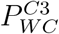 is the equilibrium WC population in *C*3. For convenience, the variables 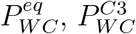 and *k_f_* are expressed below in terms of free energy differences 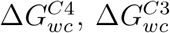 and Δ*G*_‡_ (activation free energy), respectively, using Eq. (S.5) and Eq. (S.6) at standard conditions.

First, we analyzed how 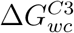 and Δ*G*_‡_ of the exoergic 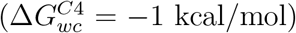 wb-WC reaction affect *η* predicted from Eq. (2) (Fig. 5B). For the range of Δ*G_wc_* and Δ*G*_‡_ in RNA duplexes in solution reported in^16^, *η* overlaps with the *in vitro* error range of G○U mismatches (~ 10^-2^ - 10^-4^)^5–9^. However, this result clearly cannot be interpreted as a validation of Eq. (2). In order to test Eq. (2), the decoding rate constants for G○U mismatches should be obtained, and Δ*G*_‡_ in the decoding site accurately estimated.

Next, we addressed the dependence of *η* on the decoding rate constants 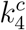 (GTPase activation) and 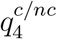 (cognate and near-cognate escape rate constants of the C4 state). These rate constants determine the deviation of the open↔closed transition in decoding from equilibrium conditions, thereby limiting the accuracy of decoding in the classical model^27,28^. Eq. (2) suggests two possible kinetic regimes of the wb-WC reaction for the case of positive 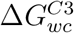. In the “fast” regime 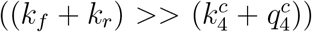, kinetics of the wb-WC reaction and 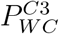 are irrelevant, and 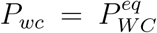. For 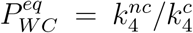 (corresponds to 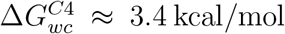), the fast regime is equivalent to the classical error *η*_0_ (Eq. (A.1)) in both 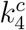 and 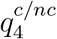 dependencies, which is confirmed by numerical calculations (Fig. 5C-D). Therefore, our model satisfies the correspondence principle^50^: it reduces to the classical induced-fit model when the wb-WC reaction is at equilibrium in the pre-chemistry step of decoding.

A more intriguing and relevant given the predicted wb-WC parameters is the “slow” regime, where the wb-WC equilibrium in *C*4 is shifted to the WC geometry (*k_f_ > k_r_*), but the kinetics of the reaction is restricted by the decoding rates. To visualize the slow regime, we chose wb-WC parameters (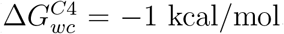, Δ*G*_‡_ = 17.8 kcal/mol) which approximately correspond to the predicted 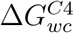 (Table 1) and experimentally observed Δ*G*_‡_ in RNA duplexes^16^. For 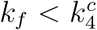, Eq. (2) predicts a virtually flat 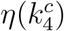 curve, as confirmed by numerical solutions, and 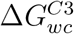 affects the offset of the curves (Fig. 5C). The reason for the flat 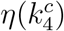 curve in the slow regime is derived in Appendix and visualized on Fig. S7: the linear approximations of *P_wc_* and 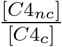 have inverse dependency on 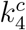, which cancels in *η*. Equilibration of the exoergic wb-WC reaction in *C*4 with unfavorable contribution to the accuracy counteracts the equilibration of the open→closed transition with favorable contribution, thereby constraining the decoding accuracy.

In 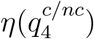, the slow kinetic regime with a very low 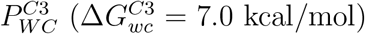 closely follows *η*_0_. However, the same kinetic regime with 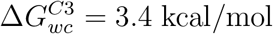 results in an almost flat curve for 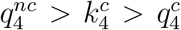 (Fig. 5D). This effect is explained by the kinetic partitioning term in 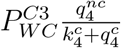, which grows proportionally with 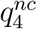 and cancels the classical equilibration process. In sum, Eq. (2), confirmed by the numerical calculations, predicts that high Δ*G*_‡_ in *C*4 causes a flat 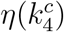 curve for 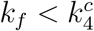, while high 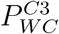 causes an almost flat 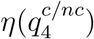 for 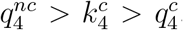. Two potentially independent parameters of the wb-WC reaction constrain the decoding accuracy of the G○U mismatch recognition in translation, which might have significant implications for the evolutionary optimization of decoding.

To explore this further, we considered a cumulative error rate of translation, *η_c_*, as a sum of *η* contributions from mismatches with slow, WC-shifted wb-WC transitions (e.g., G○U mismatches) and with “classical” *η*_0_-like behavior. In terms of our model (Fig. 5A), the latter can be interpreted as mismatches having fast endoergic transition to the WC geometry in *C*4, or mismatches whose GTPase activation rate is governed by low WC-independent rate constant (i.e. 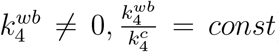), that is ignored when considering G○U mismatches. Recent studies show that the error hotspots (primarily G○U mismatches) have error rate of ~ 10^-2^ - 10^-4^ *in vitro*, and the error rate of other mismatches span a range of 10^-4^ - 10^-75-9^. Based on these findings, we approximated *η_c_* as sum of *η* of the slow kinetic regime (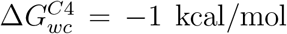, Δ*G*_‡_ = 17.8 kcal/mol, 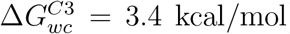) and a range of *η*_0_ curves with rescaled 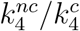 ratio that results in *η*_0_ range of 10^-3^ - 10^-8^ at the experimental values of the decoding rate constants. We calculated *η_c_* as a function of 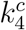 and 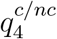. The resulting *η_c_* curves are shown as dashed lines on Fig. S8. *η_c_* is governed by *η* of the G○U mismatch for all values of 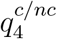, and for decreasing values of 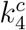.

The evolutionary optimization of the values of 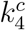 and 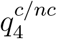 cannot be explained solely by *η* or *η_c_*, which are monotonic functions of these rate constants. To explore possible reasons for the evolutionary optimization of these rate constants to the values observed in the experiments, we looked for extrema in the gradient of *η_c_*, ∇*η_c_*. ∇*η_c_* determines the noise (i.e. standard deviation) of error rate *η_c_*, given that *η_c_* is monotonic. However, calculating the noise of 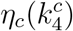 and 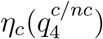 would require knowing the variance of 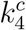 and 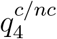. Therefore, for this illustrative purpose, we restricted to the analysis of ∇*η_c_*. Numerical gradients ∇*η_c_* were calculated for each *η_c_* curve and shown on Fig. S8 as solid lines. At *η*_0_ contribution of 10^-5^, the experimental value of 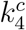 is at the local minimum of ∇*η_c_* (Fig. S8). We selected this *η*_0_ parameter to visualize the cognate rate of decoding *R_cog_*, *η_c_* and ∇*η_c_* as 2D functions of 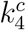 and 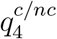 (Fig. 6). Both rate constants for a wide range do not significantly contribute to *R_cog_* (Fig. 6A, Fig. S9). The exoergic wb-WC reaction constrains *η_c_* at a plateau (Fig. 6B): for the ribosome to significantly improve the accuracy, the decoding rate needs to be reduced. In contrast, for a classical mismatch, the ribosome could reduce the error rate at least 1000-fold without significantly affecting the decoding rate (Fig. S9A).

**Figure 6:**
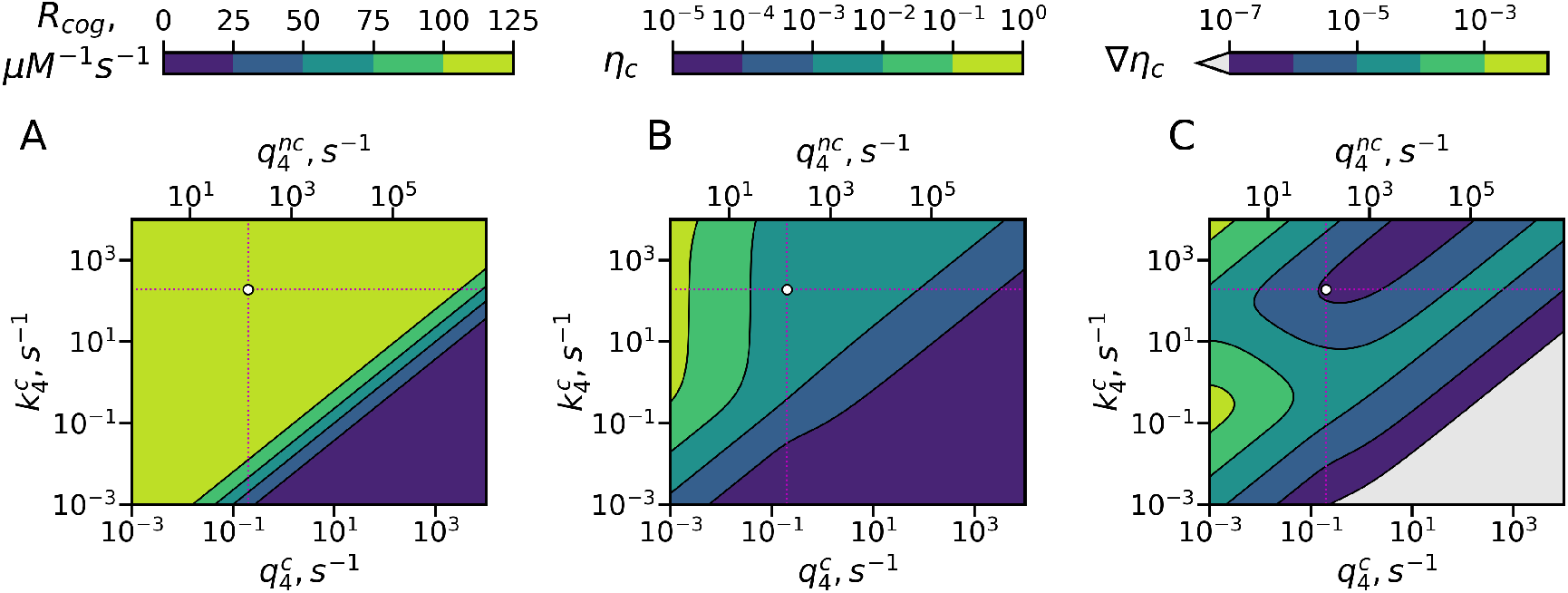
Optimization landscape of decoding constrained by *η* of G○U mismatches. The cognate rate of decoding (*R_cog_*, **A**), cumulative error rate (*η_c_*, **B**) and the gradient of cumulative error rate (∇*η_c_*, **C**) were calculated as functions of 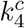 and 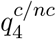. Each function was calculated as a sum of corresponding contributions from the slow regime of the wb-WC reaction (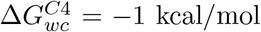, Δ*G_‡_* = 17.8 kcal/mol, 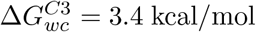) and a ‘‘classical” mismatch with rescaled 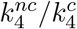 ratio that results in *η*_0_ = 10^-5^ at the experimental values of the decoding rate constants. Dotted lines and white circle at the intersection denote the experimental values of 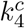 and 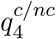.

In the cumulative approximation, the ribosome (the experimental values of decoding rate constants) is positioned in a region of local minimum of ∇*η_c_* (Fig. 6C). It illustrates the possibility of the evolutionary optimization for the minimal gradient, and thus for the minimal noise of error rate. For the case of *η* of a G○U mismatch only, this position is shifted from the minimum (Fig. S9B). Consideration of the cumulative error of decoding would likely be important to understand the evolutionary optimization of the ribosome, but requires more experimental information and more realistic approximations.

It is also worth noting that our model does not contradict the experiments with varying Mg^2+^ concentrations. In their experiments Zhang et al.^51^ reported that Mg^2+^ concentration affected *q*_2_, resulting in linear rate-accuracy trade-offs. For all considered wb-WC parameters, our model also predicts linear rate-accuracy trade-offs by varying *q*_2_ (Fig. S10).

## CONCLUDING REMARKS

Structural studies revealed stabilization of the WC geometry of the G○U base pairs in the closed ribosomal A-site^20–24^, but the mechanism of this phenomenon has been lacking a physichochemical explanation and its implications for the decoding mechanism remained unexplored. Here we addressed these problems using computational and theoretical approaches.

QM/MM umbrella sampling simulations show that closing of the rRNA residues A1492 and A1493 in the decoding site shifts the equilibrium in the wb-WC reaction to the WC geometry at the middle codon-anticodon position. This effect may partly be explained by the decreased polarity of the closed decoding site. The model of the ribosomal A-site used in our simulations is of course a simplification of the real conditions that may affect the wb-WC properties during the decoding. The major simplifications are: the model is only a part of the ribosome structure, with an outer shell that was restrained during all simulations; absent Mg^2+^ ions in the model; the “open” model differed from the “closed” model only by conformations of A1492 and A1493 residues. However, we argue that this model is able to capture the major effects of the open→closed transition on the wb-WC equilibrium. Distant structural elements and their change during the 30S domain closure are unlikely to dramatically affect the wb-WC reaction, and Mg^2+^ ions have not been reported to directly interact with the second codon-anticodon position. The role of G530, which was not affected in the open model, is however not fully explored in our study. It is remained to be investigated how the wb-WC reaction is affected in the first and third codon-anticodon positions which were not addressed in this study.

We also investigated the effects of the closed active site environment of DNA polymerases *β* and T7 on the wb-WC reaction in G○T. The wb-WC reaction on pol-*β* was exoergic, corroborating previous structural studies^17^. The wb-WC reaction equilibrium in T7-pol was not affected compared to the DNA duplex in water.

We introduced the wb-WC reaction into the classical model of initial selection in translation and derived analytical solutions for the decoding accuracy governed by the wb-WC reaction. The analytical solutions fit the numerical calculations and explain the decoding accuracy of G○U mismatch recognition in terms of the wb-WC reaction parameters in open and closed states of the A-site. This model is consistent with the WC geometry of G○U observed in the structural studies at equilibrium conditions 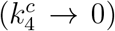^20–24^. Since a set of decoding rate constants for a G○U mismatch and the accurate 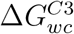 and Δ*G*_‡_ are not currently available, our model here does not serve a purpose of accurate error rate prediction. It reveals possible constraints on the codon-anticodon decoding imposed by the exoergic wb-WC tautomerization reaction: the equilibration of the reaction counteracts the equilibration of the open↔closed transition. These constraints provide a solution to the long-standing problem of seemingly suboptimal decoding mechanism pictured by the induced-fit model^26^. We speculate that the model derived here for the codon-anticodon decoding in translation can also be applicable for the base pair recognition in DNA replication. We discuss specific predictions of our model, as well as relations to the previous models and available experimental data in Supplementary Discussion. We hope that our theoretical and computational results might be a next step towards a more complete model of substrate recognition.

## Acknowledgement

We thank Prof. Andrew Torda for helpful comments on the manuscript. A.K. thanks Prof. Dmytro Hovorun for an inspiration in the problem of tautomerism of nucleobases. The work was supported by a fellowship from Deutscher Akademischer Austauschdienst (DAAD) to A.K. We are grateful for computational support by Norddeutscher Verbund für Hoch-und Höchstleistungsrechnen (HLRN) (allocation code hhc00026), and by Hummel cluster at the University of Hamburg.

## Supporting Information Available

Detailed methods, a detailed discussion of the new model, derivations of the analytical solutions of the model, Tables S1-S3 and Figures S1-S12 are available in the Supplementary Information.

## Data availability

Optimized base pair geometries, coordinate frames of the NEB trajectory, initial system topologies and coordinates in the QM/MM US simulations, as well an example setup of the QM/MM US simulations are available on Figshare: https://doi.org/10.6084/m9.figshare.c.5178074.v1

## Code availability

Python and Tcl scripts for the umbrella sampling setup, dielectric permittivity calculations, kinetic modeling and visualizations are available on GitHub: https://github.com/and-kaz/wbwc_paper

## Graphical TOC Entry

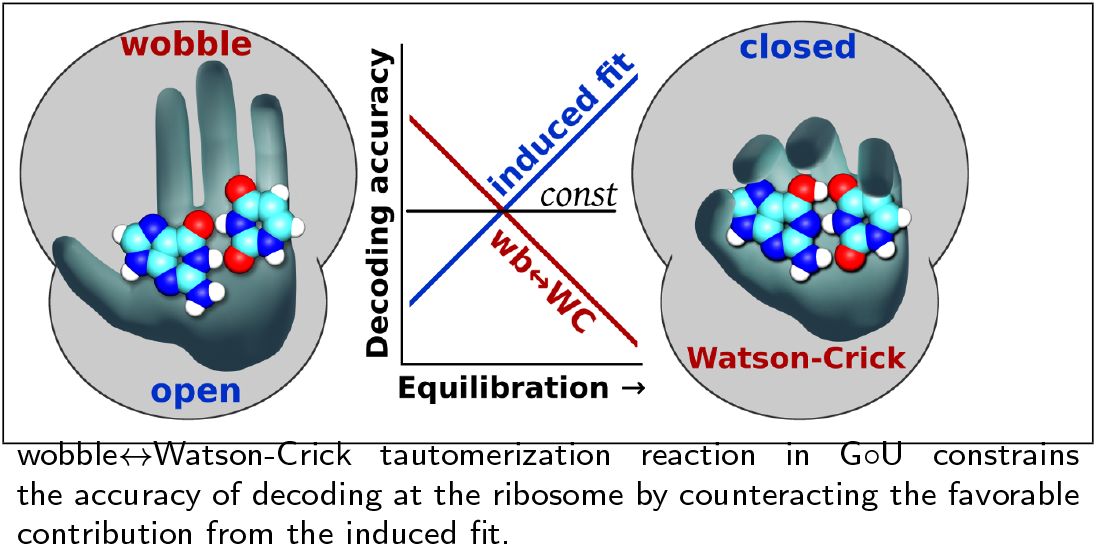

