## Supporting Information for "Tautomerization constrains the accuracy of codon-anticodon decoding"

### Supporting Information: Tautomerization constrains the accuracy of codon-anticodon decoding

### Contents

**Supplementary Methods** **S-3**

Selection of the QM level of theory for QM/MM calculations . . . . . . . . . S-3

System setup . . . . . . . . . . . . . . . . . . . . . . . . . . . . . . . . . . S-5

MD simulations . . . . . . . . . . . . . . . . . . . . . . . . . . . . . . . . S-7

Estimation of dielectric constant . . . . . . . . . . . . . . . . . . . . . . . S-8

Umbrella sampling . . . . . . . . . . . . . . . . . . . . . . . . . . . . . . S-10

Kinetic modeling . . . . . . . . . . . . . . . . . . . . . . . . . . . . . . . S-13

Analysis and visualization . . . . . . . . . . . . . . . . . . . . . . . . . S-15

**Supplementary Discussion** **S-16**

Historical perspective of the new model of substrate recognition . . . . . . . S-16

Predictions and potential refutations of the model . . . . . . . . . . . . . . S-17

Potential implications for base pair recognition in DNA replication . . . . . . S-20

**Appendix** **S-22**

**Supplementary Tables** **S-26**

**Supplementary Figures** **S-28**

**Supplementary References** **S-39**

### Supplementary Methods

#### Selection of the QM level of theory for QM/MM calculations

One of our goals was to calculate potential of mean force (PMF) of the wb-WC reaction in various molecular environments using hybrid QM/MM approach. For such calculations to converge, extensive conformational sampling is required, which would limit the use of computationally expensive high levels of theory, such as some density functional theory (DFT) methods. Therefore, we focused on faster levels of theory with acceptable accuracy. Semiempirical methods (SE) provide a great speed-accuracy ratio, but can be highly inaccurate for some systems[S1–S3]. Therefore, we first estimated the accuracy of several selected SE methods, namely PM family methods (PM3, PM6, PM6-D3 and PM7), in their ability to describe energy changes of the wb-WC reaction. Besides the gas phase energies, the optimal level of theory needed to accurately capture the effect of the implicit solvent. First, we compared total energy of the WC geometry formation ($\Delta E_{wc}$ =E(G*∘U WC) - E(G∘U wb)) calculated by selected DFT and SE methods in gas phase and in the implicit water model, using geometries optimized on $\omega$B97X-D3/def2-TZVP level. $\Delta E_{wc}$ from DLPNO-CCSD(T)/aug-cc-pVTZ was used as a reference in the calculations of absolute errors $\Delta\Delta E_{wc}$. All tested DFT (and RI-MP2) levels demonstrated small errors compared to the reference ($<$ 1.5 kcal/mol) (Table S1). All levels of theory demonstrated monotonic increase of $\Delta E_{wc}$ upon increasing relative dielectric permittivity ($\varepsilon$) of the implicit solvent model (Fig. S1).

SE levels, except for PM3, demonstrated much larger errors in $\Delta E_{wc}$ (7 – 10 kcal/mol) (Table S1). This is not unexpected, as PM methods were parameterized to accurately reproduce heat of formation ($\Delta H$), not the total energy[S4]. In order to compare $\Delta H_{wc}$ of SE and DFT methods, the latter should include zero point energy, which limits the comparison to the DFT levels with successfully optimized geometries. The sets of geometries optimized in gas phase and in implicit water model were obtained using three levels of theory: $\omega$B97X-

D3/def2-TZVP (only gas phase), B3LYP-D3BJ/def2-TZVP (no TS) and BLYP-D3BJ/def2-SVP. For each set, single-point heat of formations were calculated by the SE methods, and compared to enthalpies from the DFT methods. In this case, we did not use a single reference, but instead compared SE methods to $\Delta H_{wc}$ at the DFT level from the corresponding set of geometries. PM6 and PM6 with dispersion correction (PM6-D3) again showed large errors (4 – 9 kcal/mol) (Table S2). PM7 and PM3 showed much lower errors ($\lesssim$ 1 kcal/mol), comparable to the $\Delta H_{wc}$ differences between the selected DFT levels, and reaching the "chemical accuracy". Although PM3 method exhibited relatively high accuracy, it predicted spurious pathway of the wb-WC reaction (via ion pairs, see Fig. S2), therefore was not used for QM/MM simulations. We also compared $\Delta H_{wc}$ from PM7-optimized geometries. For this, we used two sets of DFT-optimized reference geometries (BLYP-D3BJ/def2-SVP and B3LYP-D3BJ/def2-TZVP), introduced normally distributed noise (mean = 0, SD = 0.5 Å) to the Cartesian coordinates, and subjected them to re-optimization using PM7. $\Delta H_{wc}$ from the re-optimized PM7 geometries demonstrated slightly higher, but still reasonable errors (< 2 kcal/mol) (Table S2). This indicates that not only single-point energies, but also gradients (used in geometry optimizations, and later – in QM/MM MD simulations) are reasonably accurate on the PM7 level for local minima geometries. Next, we compared SE and DFT methods based on $\Delta H_{\ddagger}$ – activation enthalpy of the wb-WC reaction. In $\Delta H_{\ddagger}$, SE methods demonstrated much higher errors (2 – 14 kcal/mol) (Table S2). PM7 performance was the best among the SE methods (2 – 8 kcal/mol errors), but still far from reaching acceptable accuracy.

Based on these benchmark calculations, we selected PM7 as a QM level of theory for all QM/MM calculations. Large errors in $\Delta H_{\ddagger}$ in PM7 do not allow using activation free energy barriers obtained from QM/MM calculations with PM7. Since a single-point and gradient calculation using PM7 takes only a fraction of a second for a base-pair-sized system, this choice of the QM method allowed us to collect enough lengths of the QM/MM MD trajectories in umbrella sampling calculations (cumulative time $\sim$ 120 ns) for the estimation of conformational sampling errors and and convergence.

All DFT, RI-MP2 and DLPNO-CCSD(T) calculations were performed in Orca 4.2.1[S5]. Tight convergence criteria were set for SCF calculations ($10^{-8}$ a.u.), as well as for geometry optimizations. Quasi-Newton optimizer using the BFGS update was used for the local minima optimizations, while Berny algorithm was used for TS optimizations. Conductor-like polarizable continuum model (CPCM) method was used for implicit solvation models[S6]. MOPAC-2016 was used for all SE calculations[S7].

#### System setup

*Benchmark systems.* Solution NMR structure of DNA dodecamer containing two wobble G∘T base pairs (PDB ID: 1BJD) was used as the initial structure for the DNA system. A heptamer centered on one of the G∘T base pairs was selected (5-CGTGACG-3, 5-CGTTACG-3). The heptamer was solvated in 50 Å x 50 Å x 50 Å box of TIP3P water[S8]. $Na^+$ and $Cl^-$ ions were added to neutralize the system and reach NaCl concentration of 0.15 M. To build the benzene system, the single G∘T base pair from the DNA system was taken after equilibration. Deoxyribose was retained, but phosphate groups were removed to have the neutral system. The base pair was solvated in 50 Å x 50 Å x 50 Å box of benzene.

*DNA-polymerases.* Although a crystal structure of pol-$\beta$ with G∘T base in WC geometry in the closed active site is available (PDB ID: 4PGX), $Mn^{2+}$ ions were used to stabilize the closed active site in this structure[S9]. Our goal was to model the effect of the native, $Mg^{2+}$-bound active site on the wb-WC reaction. Therefore, we used the crystal structure with a cognate G∘C base pair and $Mg^{2+}$ ions in the active site (PDB ID: 4KLF)[S10] as the initial structure for our pol-$\beta$ model. dCTP was replaced with dTTP. X-ray structure of T7 DNA polymerase with a cognate G∘C base pair and $Mg^{2+}$ ions in the active site and bound to thioredoxin (PDB ID: 1T7P)[S11] was used as the initial structure for our pol-T7 model. dC was replaced with dT and thioredoxin was excluded in the model. Missing protein residues in the DNA-pol models were added using Psfgen plugin in the VMD suite[S12] and were partially optimized before the equilibration protocol. Both models were solvated in TIP3P water boxes[S8], maintaining at least 16 Å from the box edges to the solute.

*A-site models.* X-ray structure of *T.thermophilus* 70S ribosome bound to tRNA$^{Thr}$ in the A-site (PDB ID: 6GSK) was used as the initial structure for all A-site models. In this structure, U∘G base pair in the second codon (AUC) - anticodon (GGU) position is solved in the WC geometry[S13]. The closed A-site model contained no manual changes in the decoding center, and the rRNA residues A1492, A1493 and G540 were in *out* conformation, surrounding the codon-anticodon helix. As a proxy of the open state of the decoding center we created abasic model, in which we deleted nucleobases of A1492, A1493 and G540 residues, leaving only their sugar-phosphate backbone. Abasic model, in contrast to a more realistic open model, alleviated the need to use harmonic restraints on these residues, which would affect dipole moment fluctuations, important in one of our analyses. Harmonic restraints to create the open state of the decoding center were used only in QM/MM umbrella sampling simulations, as described in the corresponding paragraph. Preparation of the closed and abasic models of the decoding center was performed similarly to the other studies[S14]. All residues within 35 Å radius of the center of mass (c.o.m.) of the second codon-anticodon base pair were selected for the model of the decoding site. Obtained spheres of 35 Å radius were solvated in 120 Å x 120 Å x 120 Å box of TIP3P water[S8]. $Na^+$ and $Cl^-$ ions were used to neutralize the system and reach NaCl concentration of 0.15 M. The outer $\sim$ 7 Å shell of the solute in the A-site models was restrained in all subsequent simulations with a force constant of 30 kcal mol$^{-1}$Å$^{-2}$. The inner 28 Å sphere was not restrained in the final production simulations, but was also restrained during the equilibration protocol, as described below.

All $Mg^{2+}$ ions from the ribosome X-ray structure were excluded for two reasons: 1) $Mg^{2+}$ have been shown to create artifacts when classical force fields parameters were used, thus usually requiring specialized or polarizable force fields for accurate modeling[S15,S16] and 2) in X-ray crystallography experiments, electron density peaks labeled as $Mg^{2+}$ ions may often be in fact water molecules or $Na^+$ ions[S17]. While $Mg^{2+}$ ions are undoubtedly important for ribosome structure and function[S18], their accurate modeling is out of scope in this study. Their complete exclusion may even improve the reliability of the models given that classical force fields are used, and the MD trajectories lengths are relatively short[S19]. In contrast to the ribosome crystal structures with a large number of modeled $Mg^{2+}$ ions, DNA polymerases contain 1-3 well-defined and tightly bound $Mg^{2+}$ ions, known to be essential for dNTP binding[S20]. Therefore, $Mg^{2+}$ were retained in all models of DNA polymerases.

The open A-site model in the US simulations (see below) was prepared as following. We used torsional harmonic restraints on A1492 and A1493 as in the study by Zeng et al.[S14]. By moving these harmonic restraints with a force constant of 0.04 kcal $mol^{-1}$ Å$^{-2}$, A1492 and A1493 changed their conformation from *out* to *in* and intercalated into h44 rRNA helix, characteristic of the open A-site state[S14,S21]. As the initial coordinates for the open A-site model we used frames from the end part of the US trajectories of the closed A-site model. Visual inspection of the trajectories revealed that closed→open transition happened on the timescale of approximately 10 ps.

In all models, during equilibration and all classical MD simulations, the studied G∘U(T) base pair was maintained in the WC geometry by using G* parameters from CHARMM36 force field[S22]. These parameters did not affect energies obtained in the QM/MM calculations, as the base pair was modeled with PM7 level instead.

#### MD simulations

NAMD 2.12 package was used for all MD simulations[S23]. CHARMM36 force field was used for MM part in all MD simulations[S24–S26]. Periodic boundary conditions were used in all simulations. Particle Mesh Ewald method[S27] was applied to treat electrostatic interactions and a cutoff of 12 Å used for the van der Waals interactions. Solvated and neutralized models were subjected to 1,000 steps of steepest-descent optimization of water and ions coordinates while the rest of the structure was restrained with a force constant of 50 kcal/mol/Å$^2$. Solvent and ions, as well as cell volume were equilibrated with 1-2 ns of NPT simulations with 1 fs time-step at standard conditions using Langevin thermostat and barostat, maintaining the same restraints on the solute. The obtained coordinates were used for 10,000 steps of steepest-descent optimization without any restraints apart from the outer shell in the A-site models. The optimized coordinates were used for 400 ps of gradual heating of the systems to 298 K with 1 K increment every 400 fs. Classical production simulations were conducted in NVT ensemble using Langevin thermostat at standard conditions with 2 fs time-step. SETTLE algorithm[S28] was used for rigid bonds in water while SHAKE/RATTLE algorithm[S29] was used for rigid H-containing bonds in other molecules.

#### Estimation of dielectric constant

We employed Kirkwood-Fröhlich formula (KFF) to calculate static dielectric constant $\varepsilon$ of the molecular environments surrounding the base pairs of interest. KFF relates $\varepsilon$ to dipole moment M fluctuations in a given volume[S30,S31]. We used KFF for the case of surrounding permittivity $\varepsilon_{RF} = \varepsilon$[S32]:

$$\varepsilon = \frac{3\alpha + 1 + \sqrt{9\alpha^2 + 6\alpha + 9}}{4} \tag{S.1}$$

where

$$\alpha = \frac{\langle M^2 \rangle - \langle M \rangle^2}{3\varepsilon_0 V k_B T} \tag{S.2}$$

where $\varepsilon_0$ – dielectric constant of vacuum, $k_B T$ – thermal energy, and $V$ is the volume of the probed region.

For each studied system, at least two replicas of at least 75 ns of classical MD trajectories were collected. $\varepsilon$ was measured in spheres of 5, 7, 9 and 12 Å radii centered at the base pairs of interest. For the DNA polymerases, we only focused on the G∘T base pair in the active site. For the A-site models, we calculated $\varepsilon$ in spheres surrounding each codon-anticodon base pair separately. The same replicas of MD trajectories were used for the calculations at four different radii at the three codon-anticodon positions. First 20 ns of each trajectory were excluded from the analysis. The general algorithm consisted of the following steps:

1. A center of the probed region was selected as C2 atom of the codon nucleobase at the given base pair (the template nucleobase for the DNA polymerases);

2. For a given cutoff radius, a selected trajectory was analyzed for the number of protein and nucleic acid residues present in the cutoff sphere at each MD frame; the largest set of residues was selected for the calculations, where this set was kept constant;

3. For a given cutoff radius, a selected trajectory was analyzed for the number of water molecules in the probed region. The mean number was selected for the calculations; this number was kept constant during the calculations by slight changes of the cutoff radius for water at each MD frame until the water selection converged to the needed amount of water molecules; variation of the cutoff was below 1.5 Å therefore, the cutoff radii presented on the figures reflect only the mean radii of water selection spheres;

4. The total selection at each frame consisted of the set of protein and nucleic acid residues, and the converged set of selected water molecules. The selection did not include ions, as it has been revealed previously that the dynamic contribution from ions to the static dielectric constant is relatively small and can be neglected[S33]. For this total selection, two properties were calculated: volume and dipole moment. Volume was calculated using VMD[S12] as a density map with 1.0 Å resolution; due to computational cost limitations, the density grids were calculated every 4 ns. Dipole moment was calculated by VMD[S12] using the c.o.m. of the selection as the reference point.

5. To obtain the scalar volume values, the density grids were integrated using 1.0 isovalue. Low fluctuations of the volume values from a given trajectory were confirmed, and the mean volume was used for $\varepsilon$ calculations using Eq. (S.1).

To assess the validity of our approach, we performed a benchmark calculation using a box of 42,000 TIP3P water molecules. Three replicas of approximately 30 ns were collected using classical MD in NVT ensemble at standard conditions. The described above approach was applied to this benchmark system. The center of mass of the box was used as a center of the probe spheres. The result of this benchmark calculation is shown on Fig. S3. As the Figure demonstrates, $\varepsilon$ calculated in our approach displays a size-dependence, similarly to the previous studies[S34]: $\varepsilon$ of TIP3P water converges to its experimental value of ∼80 only at approximately 20 Å radius of the probe sphere. Therefore, our approach does not allow quantitative $\varepsilon$ calculations of the relatively small regions of the decoding and active sites around the studied base pairs. We restricted our study to qualitative comparison of the closed and abasic models of the A-site, and to pol-$\beta$ and T7-pol DNA polymerases. Fig. S4 shows the cumulative mean square dipole moment fluctuation $\langle M^2\rangle - \langle M\rangle^2$ in all MD trajectories. As the Figure demonstrates, most trajectories converged to relatively constant fluctuation levels, justifying the use of KFF.

#### Umbrella sampling

*Selection of the collective variables.* wb-WC reaction involves slow motions of heavy atoms (geometry change) and fast proton transfers (PT) (Fig. 1A), making it impossible to describe with a 1D collective variable (CV). Therefore, the geometry change and PTs were described with two separate CVs. To describe the geometry change, we used path collective variable (pathCV). PathCV requires a set of structures (images) describing the process and used as a reference[S35]. In pathCV, the position of a given coordinate frame on the path ($s$) is calculated as:

$$s = \frac{\sum_{i=1}^{N} i \exp(-\lambda R[X - X_i])}{\sum_{i=1}^{N} \exp(-\lambda R[X - X_i])} \tag{S.3}$$

and the distance from the path (z):

$$z = -\frac{1}{\lambda} \ln \left[ \sum_{i=1}^{N} \exp(-\lambda R[X - X_i]) \right] \tag{S.4}$$

where $R[X - X_i]$ is a distance metric, describing the distance from a given frame to the path image $i$. As a distance metric, we used RMSD as implemented in colvars module of

NAMD 2.12[S36]. $\lambda$ is a parameter that can be tuned for optimal performance of pathCV. We used $\lambda$ value of 300 throughout all pathCV calculations.

To obtain the reference path, we optimized the full wb-WC reaction in G∘U *in vacuo* using nudged elastic band method (NEB). We used NEB-TS implementation in Orca 4.2.1[S5], which is a combination of climbing-image NEB (CI-NEB)[S37] and eigenvector-following optimization of a TS guess. NEB-TS was performed on B3LYP-D3BJ/def2-TZVP level of theory using the default spring force constant of 0.1 Eh Bohr$^{-2}$ and tight criteria for SCF convergence. The optimized path contained 34 frames. Its potential energy profile is shown on Fig. 1B. To create the set of images for pathCV, the double-proton-transfer (DPT) part of the NEB path (frames 26 to 34) was excluded, as it did not contain the geometry changes. Only ring atoms of the nucleobases were included into pathCV calculations. PTs in the wb-WC reaction were described as a distance difference $hb$ = d(O6-H3) - d(N1-H1). Characterization of the wb-WC reaction in 2D CV space ($s$; $hb$) clearly distinguished three local minima and the TS on the reference NEB path (Fig. 1C). Although the previous computational studies revealed only G∘U wb → G∘U* WC path in the wb-WC reaction[S38], the possibility of the alternative path G∘U wb → G*∘U WC, or a bifurcation leading to both products could not be excluded. To verify the TS from the NEB calculations, and to exclude at least the post-TS bifurcation, we performed committor analysis in gas phase on BLYP-D3BJ/def2-SVP. Approximately 50 Born-Oppenheimer MD simulations were started from the TS with randomly initialized velocities matching 298 K using Berendsen thermostat with 2 fs period. Simulations were performed for 200 fs with 1 fs time step. Committor analysis revealed roughly equal partition between reactant (wb) and product (G∘U* WC), suggesting validity of the TS (Fig. 1C). No trajectories led to G*∘U WC, which enables excluding a post-TS bifurcation in this reaction (Fig. 1C). Therefore, selected reference path and CVs accurately describe the wb-WC reaction.

*Setup of the US calculations.* To calculate potential of mean force (PMF) of the wb-WC reaction in the selected models, we applied umbrella sampling (US) simulations. US allows generating a biased ensemble in a series of windows using specified CV-derived biases[S39]. Using weighted histogram analysis method (WHAM), applied to the set of biased distributions in windows, the unbiased PMF can be calculated[S39]. We used 34 frames from the NEB path as the initial base pair coordinates of the US windows. For each studied system, a frame from the end part of the corresponding classical MD trajectory was selected as the initial system coordinates for US simulations. To prepare the initial US windows, coordinates of the studied G∘U(T) base pair in the initial system were changed to the pre-aligned coordinates of each NEB frame. Thus, 34 initial US windows in each simulated system differed only by the base pair coordinates. $z$ was not used as a CV for PMF calculation. Instead, in all US simulations, a half-harmonic boundary potential was added at $z$ value of 0.15 with a force constant of 100 kcal/mol/Å$^2$. This prevented simulations from visiting largely out-of-plane or shifted base pair conformations, where $s$ would not be well-defined. At the same time, using the boundary potential instead of restraining $z$ at 0 allowed us to ignore $z$ in WHAM calculations.

To improve sampling in the vicinity of the minima while still covering the full wb-WC path, we used two layers of US windows with different force constants applied to the chosen CVs $s$ and $hb$. The more rigid layer with the force constant of 50 kcal/mol/Å$^2$ applied to both $s$ and $hb$ covered all US frames in the TS region and selected frames in the vicinity of the minima. The more flexible layer with the force constant of 5 kcal/mol/Å$^2$ only covered frames in the vicinity of the minima. The two layers together contained 48 US windows per each system.

*QM/MM US simulations.* All US simulations were performed in hybrid quantum-mechanical/ molecular-mechanical (QM/MM) scheme. PM7[S4] in MOPAC2016[S7] was used for the QM region and CHARMM36 force field[S24–S26] in NAMD2.12[S40] was used for the MM region. In all systems, the QM region comprised only the nucleobases in the base pair of interest (26-29 atoms), with the QM/MM interface placed at the glycosidic bonds. The hydrogen link-atom approach with charge shift scheme was used to treat the link atoms. Tight

SCF convergence criteria ($10^{-8}$ kcal/mol) were used in PM7 calculations in MOPAC. It is worth emphasizing that in the MOPAC2016/NAMD inteface, NAMD reads heats of formation from MOPAC[S40]. Each US window was minimized with 100 steps of steepest-descent before the start of the production simulations. All US simulations were performed in NVT ensemble using Langevin thermostat at standard conditions. Integration step was 0.2 fs, and trajectories were collected every 40 fs. First 12 ps of the US trajectories were discarded from the analysis. Each US window was simulated for at least 100 ps up to 400 ps, resulting in the cumulative ~120 ns of the QM/MM simulations. In each system, some (random) US trajectories lagged behind the rest of the trajectories, as they were running individually on the computing cluster. These trajectories were excluded from the WHAM calculations.

*PMF calculations.* wham-2D code by Alan Grossfield was used for all WHAM calculations. US trajectories were divided into batches of approximately 20 ps, and PMF was calculated from each batch separately to monitor the convergence. The grid for WHAM calculations was [0;1] with 0.025 step for $s$ and [-1.8;1.8] with 0.05 Å step for $hb$. Empty bins were filtered out. The grid was divided into regions corresponding to G∘U wb, G∘U* WC and G*∘U WC basins. $\Delta G$ of each of the three states was calculated as local PMF minima in each basin. For each batch, $\Delta G_{wc}$ was calculated as $\Delta G$(G*∘U WC) - $\Delta G$(G∘U wb). Visual inspection of the $\Delta G_{wc}$ convergence allowed to arbitrary assign converged regions of the trajectories (Fig. 4B-D). These regions were then treated as a single batch in each trajectory, PMFs from which are shown on Fig. 4A and Fig. S5. MEPSA was used to calculate a minimal free energy path from PMF[S41].

#### Kinetic modeling

*Rate constants.* Decoding rate constants for a codon-anticodon combination with a G∘U mismatch in the first position are avialable only for the low-fidelity conditions[S42], corresponding to unrealistically low accuracy of initial selection compared to the more recent experiments[S43–S47]. Therefore, we selected the set of rate constants corresponding to the

A∘C mismatch in the first codon-anticodon position measured at high-fidelity conditions[S48] (Table S3). C2↔C3 transition (codon reading) is very rapid, which prevents estimation of its rate constants[S49]. Therefore, we used arbitrary rate constants for this transition, namely $k_2, q_3^c$ and $q_3^{nc}$. In order to maintain at least some consistency with the previous studies, we calculated these rate constants from a free energy diagram in Pavlov and Ehrenberg[S50]. However, the low free energy barriers in this diagram result in extremely high rate constants ($\sim 10^9$ s$^{-1}$) that would prohibit the use of numerical calculations. Therefore, we uniformly scaled these rate constants down to values acceptable for the numerical calculations, while keeping the values high enough to not have significant rate-limiting effects (Table S3).

All equilibrium populations were calculated from the free energy changes $\Delta G$ according to Boltzmann population at standard conditions, or equivalently, from forward and reverse rate constants:

$$P_{eq} = \frac{\exp \frac{-\Delta G}{RT}}{\exp \frac{-\Delta G}{RT} + 1} = \frac{\frac{k_f}{k_r}}{\frac{k_f}{k_r} + 1} \tag{S.5}$$

where $RT$ is the thermal energy.

Rate constants were calculated from the free energy of activation $\Delta G_{\ddagger}$ according to Eyring equation with transmission coefficient 1 at standard conditions:

$$k = \frac{k_B T}{h} \exp \frac{-\Delta G_{\ddagger}}{RT} \tag{S.6}$$

where $k_B$ is Boltzmann's constant and $h$ is Planck's constant.

*Numerical calculations.* Numerical solutions of the kinetic systems of decoding were obtained by numerical integration of ordinary differential equations (ODE) in Python 3.6, describing the kinetic model on Fig. 5A. The same rate constants were used for both analytical solutions and numerical calculations. As the initial conditions, we used $[R_1] = 50\mu M$, $[T_c] = 1\mu M$, $[T_{nc}] = 1\mu M$, and zero concentrations for the rest of the states. A close to constant concentrations of $T_c$ and $T_{nc}$ were maintained by using rapid zero-order formation reactions and first-order degradation reactions, rate constants of which were set to result in the desired concentrations of $T_c$ and $T_{nc}$. Rapid equilibrium approximation for the wb-WC reaction in states C2 and C3 was maintained by using very low $\Delta G_{\ddagger}$ in these states ($6-8$ kcal/mol). Numerical integration at each point in the space of rate constants was performed for 800 s with 2 $\mu$s step ($4 \cdot 10^8$ steps). The concentrations from the last step were used as steady-state concentrations. $\eta(ODE)$ was calculated as $\frac{[P_W]}{[P_R]}$. $P_{wc}(ODE)$ was calculated as $\frac{[C4_{nc}^{WC}]}{[C4_{nc}^{WC}]+[C4_{nc}^{wb}]}$.

Derivations of analytical solutions are described in Appendix.

#### Analysis and visualization

All data analysis and visualization was performed in Python 3.6 using NumPy[S51] and Matplotlib[S52], unless specified otherwise. Visualization of molecular structures were performed in VMD[S12] and Chemcraft.

### Supplementary Discussion

#### Historical perspective of the new model of substrate recognition

The historical development of substrate recognition models reflects the increasing appreciation of the flexibility in ribo-/enzyme–substrate binding. The earliest ”lock-and-key” model considered both partners as rigid bodies with a steric complementarity to each other[S53]. Later, the inability of the lock-and-key model to explain substrate specificity of some enzymes led Koshland to propose the ”induced-fit” model[S54]. This model posits that a cognate substrate induces changes in the enzyme's active site, required for the enzyme's catalytic activity. The ”conformational selection” model, proposed later, suggests that the ”active” conformational states of an enzyme exist even in the absence of a cognate substrate, and are only stabilized by its binding[S55]. Both induced-fit and conformational selection models focus only on the enzyme flexibility, and continue to assume a rigid substrate. While such assumption is likely valid for a majority of small-molecule substrates, it arguably oversimplifies the base pair recognition. A possible indication of the inability of the rigid-substrate models to explain the mechanism of base pair recognition came from structural studies of DNA polymerases[S9,S56] and ribosome[S13,S57–S60], revealing the WC-like geometries of mismatches in the closed states of the decoding/active sites. Theoretical predictions of the induced-fit/conformational-selection models highlight this issue from another angle. Based on the mechanism of accuracy facilitation via open→closed transition, predicted by these models, the ribosome seems to be in a suboptimal space of decoding rate constants[S61], likely indicating incomplete models.

In this study we developed a model which can provide solutions to both structural and theoretical challenges unresolved by the induced-fit model. This new model builds on the induced-fit model, but allows considering flexible substrates that change *during* the decoding. On Fig. S11 we illustrate the evolution of the substrate recognition models using simplified schemes of the three discussed models. The scheme also illustrates the way the models can be reduced to the preceding ones. In accordance with the correspondence principle by Bohr[S62], our model reduces to the induced-fit/conformational-selection model upon equilibrium in the substrate at the last pre-chemistry step. Similarly, the induced-fit/conformational-selection model reduces to the lock-and-key model upon equilibrium in ribo-/enzyme.

In contrast to the previous theoretical studies proposing alternative explanations for the evolutionary optimization of decoding[S63,S64], our model has a well-defined predictive power. Below we discuss specific predictions and potential refutations of the new model.

#### Predictions and potential refutations of the model

*Constrained error rate.* The new model predicts a vastly different dependence of the error rate $\eta$ on $q_4^{c/nc}$ and $k_4^c$ compared to the classical model. In the classical model the dependence of $\eta_0$ on $k_4^{c/nc}$ is governed by the equilibration of cognate to near-cognate ratio of decoding state populations. In our model, this process is counteracted by the equilibration of the wb-WC reaction in C4, leading to the virtually flat $\eta(k_4^c)$ for a range of $k_4^c$ where the wb-WC reaction is out of equilibrium. Dependence of the classical $\eta_0$ on $q_4^{c/nc}$ is symmetric to $\eta_0(k_4^c)$, as it is also governed by the same equilibration process. In our model, the equilibrium WC population in C3 contributes to the nearly flat region in $\eta(q_4^{c/nc})$ for $q_4^{nc} > k_4^c > q_4^c$, as its kinetic partitioning into C4 grows proportionally with $q_4^{nc}$, thus counteracting the "classical" equilibration process.

To the best of our knowledge, currently available studies of rate-accuracy trade-offs do not confirm or disprove this prediction of our model.

Thompson and Karim[S65] used a slowly-hydrolyzable GTP analog GTP[$\gamma$S] to study the binding of cognate (tRNA$^{Phe}$) and non-cognate (tRNA$^{Leu}_{CAG}$) ternary complexes to the poly(U)-programmed ribosomes. They concluded that the observed $K_d^c/K_d^{nc}$ ratio, lower than would have been expected in experiments with GTP, was due to the reduced rate of GTP[$\gamma$S] hydrolysis, and therefore the decoding was close to equilibrium conditions[S65]. One of the features of this study that makes it not a refutation of our prediction is the use of non-cognate tRNA with two mismatches (U∘G in the first and U∘C in the third codon-anticodon positions).

A study by Zeidler et al.[S66] concluded that H85Q variant of EF-Tu decreased the error of Leu vs Phe misincorporation from bulk tRNA on poly(U)-programmed *T. thermophilus* ribosomes at 65 °C due to a decreased rate of GTP hydrolysis by this variant. However, in their study the effect of the H85Q variant on tRNA binding was higher than on the GTP hydrolysis[S66], therefore the source of the decreased error could not be clearly determined. Moreover, the contribution of individual mismatches could not be estimated.

Although not available yet, testing $\eta(k_4^c)$ dependence seems feasible by e.g. GTP analogs with lower or higher hydrolysis rate by EF-Tu. Testing the prediction of $\eta$ dependence on increasing $(q_4^{c/nc})$ might be more complicated, as it would require selective destabilization of the closed A-site state. The error-restrictive mutations in ribosomal protein S12 in the study by Zaher and Green[S67] did not significantly affect $(q_4^{c/nc})$, but rather the steps following the initial selection.

It is important to emphasize that for $q_4^{nc} < k_4^c$ and for a reasonable range of $\Delta G_{\ddagger}$, our model predicts similar behavior of $\eta(q_4^{c/nc})$ as in the classical model (Fig. 5D), thus not contradicting the experiments with aminoglycosides[S68] and ribosome ambiguity mutations[S69].

*Constrained near-cognate GTPase activation rate constant.* In the linear approximation, for $k_r << k_f < k_4^c$ we can derive an equality (see Appendix):

$$k_4^{nc} = k_f + q_4^{nc} P_{WC}^{C3} \tag{S.7}$$

Eq. (S.7) suggests that the apparent near-cognate rate constant of GTPase activation $k_4^{nc}$ is defined by the parameters of the wb-WC reaction for a biologically-relevant range of $k_4^c$. $k_4^{nc}$ of a codon-anticodon combination with a G∘U mismatch is yet to be accurately measured. In DNA replication, the similarity between $k_{pol}^{incorrect}$ and $k_f$ values was already noted by Kimsey et al.[S70], but remained unexplained within their numerical kinetic modeling approach.

*Symmetry in the rate constants of the open↔closed transition.* A theoretical study by Savir and Tlusty[S63] attributed the "optimal" values of decoding rate constants $q_4^{c/nc}$ and $k_4^{c/nc}$, in which $q_4^{nc}$ and $k_4^c$ are similar, to a symmetry in a fitness solution that optimizes both rate and accuracy. Since neither of these rate constants for a broad range significantly contributes to the cognate rate of even the initial selection (given the *in vitro* rate constants values), it remained unclear how exactly such solution optimizes fitness. Our model provides an alternative explanation. In our model the apparent similarity between $q_4^{nc}$ and $k_4^c$ can be explained with the kinetic partitioning term $\frac{q_4^{nc}}{k_4^c+q_4^c}$ in $P_{wc}^0$. For $q_4^{nc} > k_4^c$, and assuming $q_4^c << k_4^c$, contributions of $P_{wc}^0$ to the error rate will exceed contributions from the slow wb-WC reaction in C4 (i.e. $k_f$), and mostly cancel the classical equilibration, thus having virtually no advantage to the accuracy of decoding. For $q_4^{nc} < k_4^c$, the error is dominated by $k_f$ in C4 and increases similarly to the classical model. It is possible to suggest that decoding in translation was optimized under the constraints of two independent wb-WC parameters $P_{wc}^{C3}$ and $k_f$, resulting in an "optimal" solution where both parameters have almost equal contribution to the error rate. Although our model can justify the ribosome position at $q_4^{nc} \approx k_4^c$, a rigorous explanation would still require a consideration of additional variables, since $q_4^{nc} \approx k_4^c$ is not a local minimum of $\eta$.

*Implications for proof-reading.* Our model of the wb-WC reaction effects on decoding accuracy was restricted to the initial selection, but it does not contradict the concept of proof-reading in translation. Recent cryo-EM studies demonstrate that in post-GTP-hydrolysis states the decoding site resets back to the open state[S71], which could potentially reset the equilibrium in the wb-WC reaction back to the wobble geometry. However, proof-reading of G∘U mismatches in translation is yet to be directly demonstrated. The excess of GTP hydrolysis compared to the peptide bond formation, used in quantifying proof-reading and implicitly assumed to arise from dissociation of the near-cognate ternary complexes after the initial selection[S72], may also result from multiple rounds of EF-Tu rebinding and GTP hydrolysis with the same aa-tRNA in the A-site[S73].

#### Potential implications for base pair recognition in DNA replication

In this study we focused mainly on the role of the wb-WC reaction in codon-anticodon decoding, but its role in base pair recognition during DNA replication was also relevant to us. Our QM/MM US calculations showed exoergic wb-WC reaction in G∘T base pair in the closed active site of pol-$\beta$, in accordance with previous structural studies[S9]. In contrast, the $\Delta G_{wc}$ of G∘T in the closed active site of T7 polymerase was close to its value in DNA duplex in water. The polarity of their active sites was not drastically different, suggesting a major role of tighter steric constraints on the base pair geometry in pol-$\beta$ active site. Both DNA polymerases employ similar recognition mechanisms, which is also fundamentally similar to codon-anticodon recognition in translation (i.e. the presence of the open→closed transition of the active/decoding site[S74]). Therefore, it is reasonable to suggest that some DNA polymerases can be plagued by the wb-WC reaction in G∘T in a similar way as our model predicts it for the ribosome. We speculate that dependence of error rate on the rate constants of the open→closed transition in pol-$\beta$ would also display flat regions. It is also intriguing to speculate that the potentially reduced steric constraints on the WC geometry in the active site of T7-pol (which is in line with the absence of structural studies showing the WC geometry of G∘T in the active site of this DNA polymerase) could be an adaptation to allow T7-pol to operate in close to equilibrium conditions of G∘T recognition with respect to the wb-WC kinetics. Previous theoretical studies have suggested that this DNA polymerase operates in the energetic regime of recognition, as opposed to the kinetic regime in another DNA polymerase, pol-$\gamma$[S75].

*Relation of our model and a numerical kinetic model from Kimsey et al.[S70].* The recent study by Kimsey et al.[S70] may seem as a refutation of our model applied to DNA polymerases. Since Al-Hashimi and coauthors[S70] could predict misincorporation rates of several DNA polymerases from the equilibrium WC geometry populations in solution, it may indicate that no stabilization of the WC geometry happens in the active sites of any of the DNA polymerases they studied, or such stabilization does not affect the error rate.

However, this experimental observation does not contradict our model. In the numerical kinetic modeling approach employed by Kimsey et al.[S70], the wb-WC reaction was modeled only in the open state of the active site, using the experimentally measured $\Delta G_{wc}$ in water. Continuing our analogy with the ribosome, it is reasonable to suggest that the environment of the open active site of DNA polymerases does not significantly perturb the WC population compared to water solution. The linearity of the model by Kimsey et al.[S70] results in $P_{wc}^{closed} = P_{wc}^{open}$ (i.e. the wb-WC reaction in the closed state is absent, and the open→closed kinetic partition term equals 1). In our model, in the vicinity of experimental $\Delta G_{\ddagger}$ and $\Delta G_{wc}$ values, and for the used values of the decoding rate constants, Eq. (2) predicts comparable contributions to $\eta$ from $P_{wc}^{0}$ and from the slow wb-WC reaction in C4 ($k_f$) (Fig. 5B). In fact, as discussed above, we speculate that the similarity of the contributions from $P_{wc}^{0}$ and $k_f$ could be a result of the evolutionary optimization under the constraints of these two independent parameters. Therefore, the numerical kinetic model by Kimsey et al.[S70] (implicitly) predicted $P_{wc}^{open}$ from the experimental data, which can still be a good approximation to $P_{wc}^{closed}$ given that $P_{wc}^{open} \approx \frac{k_f}{q^{nc}_{closed\rightarrow open}}$. If in some DNA polymerases the equilibrium of the wb-WC reaction is not significantly affected by the closed active site environment, like our QM/MM calculations predict it for T7-pol (Fig. 4), $P_{wc}^{closed} \approx P_{wc}^{open}$ would be an even more realistic approximation. Therefore, the study by Kimsey et al.[S70] is neither a confirmation, nor a refutation of our model. Since $\Delta G_{wc}$ and $\Delta G_{\ddagger}$ of the wb-WC reaction measured by Kimsey et al.[S70] in duplexes in solution were correlated (Fig. S12), it can be challenging to experimentally distinguish predictions of our model from the interpretations by Kimsey et al.[S70].

### Appendix

**Derivation of Eq. (2).** Nomenclature of the states and decoding rate constants is explained in details elsewhere[S50]. The scheme of the ”classical” initial selection in decoding is shown on Fig. S6. In the Michaelis-Menten formulation, the error of decoding in translation $\eta_0$:

$$\eta_0 = \frac{R^{nc}}{R^c} = \frac{(k_{cat}/K_m)^{nc}}{(k_{cat}/K_m)^c} = \frac{k_4^{nc}[C4_{nc}]}{k_4^c[C4_c]} \tag{A.1}$$

where $R^i$ is the rate of decoding, $k^i_{cat}$ is the catalytic rate constant, $K^i_m$ is the Michaelis-Menten constant, $[C4_i]$ is the steady-state concentration of $C4$ state, and $k^i_4$ is the rate constant of GTPase activation, for $i = c$ (cognate), $nc$ (near-cognate).

According to Pavlov and Ehrenberg[S50], $R^i$ can be expressed in terms of rate constants as following:

$$R^i = \frac{k_1}{1 + a^i_2(1 + a^i_3(1 + a^i_4))} \tag{A.2}$$

where $a^i_2 = q_2/k_2$, $a^i_3 = q^i_3/k_3$, $a^i_4 = q^i_4/k^i_4$.

Now let us consider the decoding scheme on Fig. 5A. $R^{nc}$ can be expressed as following :

$$k_4^{nc}[C4_{nc}] = k_4^{wb}[C4^{wb}_{nc}] + k_4^{wc}[C4^{wc}_{nc}] \tag{A.3}$$

Given $k_4^{wb} = 0$ *and* $k_4^{wc} = k_4^c$ (see main text) and from Eq. (A.3), we obtain Eq. (1). From Eq. (A.1) and Eq. (1), the error from the wb-WC reaction in the $C4$ state $\eta$:

$$\eta = \frac{[C4_{nc}]}{[C4_c]} P_{wc} \tag{A.4}$$

Now, we can write $P_{wc}$ (WC population in state $C4_{nc}$) in terms of forward and reverse rate constants of the tautomerization reaction $(k_f, k_r)$ using equation for product concentration in reversible first order chemical reaction at time $\tau$:

$$P_{WC} = P_{wc}^{eq} + (P_{wc}^{0} - P_{wc}^{eq}) \exp\left(-(k_f + k_r)\tau\right) \tag{A.5}$$

where $P_{wc}^{eq}$ is the equilibrium WC population in $C4_{nc}$ for a given $(k_f, k_r)$, and $P_{wc}^{0}$ is the initial WC population in $C4_{nc}$. The meaning of $P_{wc}^{0}$ is the contribution of $P_{wc}^{C3}$ to $P_{wc}$. Such contribution is affected by the relative forward and reverse $C3 \leftrightarrow C4$ rates between wb and WC "branches". Therefore, $P_{wc}^{0}$ is the WC population in $C3$ state, kinetically partitioned into $C4$ state:

$$P_{wc}^{0} = P_{wc}^{C3} \frac{K_{C3\to C4}^{wc}}{K_{C3\to C4}^{wb}} = P_{wc}^{C3} \frac{q_4^{nc}}{k_4^c + q_4^c} \tag{A.6}$$

where $K_{C3\to C4}^{wc} = \frac{k_3}{k_4^c + q_4^c}$ and $K_{C3\to C4}^{wb} = \frac{k_3}{q_4^{nc}}$.

$\tau$ has a meaning of a lifetime over which the product can form. The residence time of $C4_{nc}^{wc}$ state (see Fig. 5A):

$$\tau = \frac{1}{k_4^c + q_4^c} \tag{A.7}$$

From Eq. (A.4), Eq. (A.5) and Eq. (A.7) we obtain Eq. (2). When calculating $\eta$ by equation Eq. (2) using Eq. (A.2), $k_4^{nc}$ should be substituted with $P_{wc} k_4^c$ according to Eq. (1).

**Linear approximation of Eq. (2).** Linear approximations were used to visualize the cancelling of $\eta(k_4^c)$ dependence in the slow kinetic regime of the wb-WC reaction, and to derive Eq. (S.7). For the linear approximation, we assumed the slow kinetic regime of the reaction. With this assumption, $P_{wc}^{eq} \approx 1$ since $\Delta G_{wc}$ is negative. Also, $k_r$ in the numerator of the exponential term in Eq. (2) can be omitted since $k_f >> k_r$. To simplify the solution even further, we assumed $q_4^c << k_4^c$, and thus we can approximate $\tau = 1/k_4^c$ and $P_{wc}^{0} = P_{wc}^{C3} \frac{q_4^{nc}}{k_4^c}$. Under these assumptions we can write:

$$P_{wc} = 1 + (P_{wc}^{0} - 1) \exp\left(-\frac{k_f}{k_4^c}\right) \tag{A.8}$$

To derive the linear approximation from Eq. (A.8) we also assume $k_f < k_4^c$. This allows to neglect higher-order terms in the Taylor expansion of the exponential term in Eq. (A.8). Thus, the linear approximation of Eq. (A.5):

$$P_{wc}^L = 1 + (P_{wc}^0 - 1)\left(1 - \frac{k_f}{k_4^c}\right) = \frac{k_f + q_4^{nc} P_{wc}^{C3}}{k_4^c} + \frac{k_f q_4^{nc} P_{wc}^{C3}}{(k_4^c)^2} \approx \frac{k_f + q_4^{nc} P_{wc}^{C3}}{k_4^c} \tag{A.9}$$

Fig. S7 demonstrates the validity of the linear approximation Eq. (A.9), as it matches non-approximated Eq. (A.5) and numerical calculations for the biologically-relevant region of $k_4^c$. By expressing $P_{wc}^L$ in terms of $k_4^c$ and $k_4^{nc}$ according to Eq. (1), we obtain Eq. (S.7).

With Eq. (A.9) we expressed $P_{wc}^L$ as a linear function of $k_4^c$. However, the cancelling of $k_4^c$ in $\eta$ of the slow kinetic regime of the wb-WC reaction is still not immediately evident. To clearly observe this, we also needed to simplify $\frac{[C4_{nc}]}{[C4_c]}$ (designated as $\frac{1}{D}$ below). To do this, we first considered equilibration of $D$ as a function of $k_4^c$. At $k_4^c \to 0$, $D$ approaches its equilibrium value $D_{max}$. $D_{max}$ is the "intrinsic selectivity" from Pavlov and Ehrenberg[S50], multiplied by $\frac{k_4^{nc}}{k_4^c}$:

$$D_{max} = \frac{a_3^{nc} a_4^{nc}}{a_3^c a_4^c} \frac{k_4^{nc}}{k_4^c} = \frac{q_3^{nc} q_4^{nc}}{q_3^c q_4^c} \tag{A.10}$$

At $k_4^c \to \infty$, $D$ approaches its minimal value $D_{min}$. The exact form of $D_{min}$ is irrelevant below, as $D_{min} << D_{max}$. Using these two variables as initial and equilibrium concentrations, we can describe the process of $D$ equilibration using the equation for a first-order reversible reaction:

$$D = D_{max} + (D_{min} - D_{max}) \exp\left(-\frac{Q}{K}\right) \approx D_{max} - D_{max} \exp\left(-\frac{Q}{K}\right) \tag{A.11}$$

where $Q$ and $K$ are total reverse and forward rates of decoding. The equilibration of $D$ is limited by the cognate decoding rates[S50]. Therefore, we can approximate $Q$ and $K$ from the cognate rate constants only:

$$Q = \prod q_i^c = q_2 q_3^c q_4^c \tag{A.12}$$

and

$$K = \prod k_i^c = k_2 k_3 k_4^c \tag{A.13}$$

Since we are interested in the non-equilibrium region where $K > Q$, the ratio in the exponential term in Eq. (A.11) allows to apply the linear approximation to the Taylor expansion of the exponential, similarly as performed above to obtain Eq. (A.9):

$$D^L = D_{max} - D_{max}\left(1 - \frac{Q}{K}\right) = \frac{QD_{max}}{K} \tag{A.14}$$

Using Eq. (A.10), Eq. (A.12) and Eq. (A.13), from Eq. (A.14) we obtain the linear approximation of $\frac{1}{D}$:

$$\left(\frac{1}{D}\right)^L = \frac{K}{QD_{max}} = \frac{k_2 k_3 k_4^c}{q_2 q_3^{nc} q_4^{nc}} \tag{A.15}$$

Eq. (A.15) is a good approximation to $\frac{[C4_{nc}]}{[C4_c]}$ in the non-equilibrium region, as demonstrated on Fig. S7.

From Eq. (A.15) and Eq. (A.9) we obtain the linear approximation of $\eta$ for the slow kinetic regime of the wb-WC reaction at non-equilibrium conditions:

$$\eta^L = \left(\frac{1}{D}\right)^L P_{wc}^L = \frac{k_2 k_3}{q_2 q_3^{nc}}\left(\frac{k_f}{q_4^{nc}} + P_{wc}^{C3}\right) \tag{A.16}$$

which is indeed independent of $k_4^c$. Fig. S7 demonstrates the validity of Eq. (A.16).

### Supplementary Tables

**Supplementary Table S1:** Benchmark calculations of total energy of the WC geometry formation

| Method | $\Delta E_{wc}^{gas}$ [a] | $\Delta E_{wc}^{water}$ [b] | $\Delta\Delta E_{wc}^{gas}$ [c] | $\Delta\Delta E_{wc}^{water}$ [d] |
|---|---|---|---|---|
| DLPNO-CCSD(T)/aug-cc-pVTZ | -1.98 | 3.17 | 0.0 | 0.0 |
| RI-MP2/def2-TZVP | -1.93 | 3.0 | 0.05 | 0.17 |
| $\omega$B97X-D3/def2-TZVP | -1.11 | 4.53 | 0.87 | 1.35 |
| $\omega$B97X-V/def2-TZVP | -1.21 | 4.38 | 0.77 | 1.2 |
| B97M-V/def2-TZVP | -2.05 | 3.55 | 0.07 | 0.38 |
| B3LYP-D3BJ/def2-TZVP | -1.65 | 3.94 | 0.33 | 0.77 |
| M06-2X/6-31+G(d,p) | -3.31 | 2.18 | 1.33 | 0.99 |
| BLYP-D3BJ/def2-SVP | -1.15 | 3.73 | 0.83 | 0.56 |
| b97-3c | -1.5 | 4.21 | 0.48 | 1.03 |
| PBEh-3c | -2.55 | 2.72 | 0.57 | 0.45 |
| PM3 | -1.26 | 3.8 | 0.72 | 0.62 |
| PM6 | 6.6 | 12.9 | 8.58 | 9.73 |
| PM6-D3 | 6.6 | 12.9 | 8.58 | 9.73 |
| PM7 | 3.74 | 10.11 | 5.72 | 6.93 |

[a] Total energy of the WC formation in gas phase (E(G*∘U WC) - E(G∘U wb)); [b] Total energy of the WC formation in implicit water model; [c] Absolute error in gas phase; [d] Absolute error in implicit water model; All energies are given in kcal/mol. All calculations were performed on base pair geometries optimized at $\omega$B97X-D3/def2-TZVP level of theory in gas phase.

**Supplementary Table S2:** Benchmark calculations of enthalpy of the WC geometry formation and activation enthalpy

| Method | $\Delta H_{wc}^{gas}$ [a] | $\Delta H_{wc}^{water}$ [b] | $\Delta H_{\ddagger}^{gas}$ [c] | $\Delta H_{\ddagger}^{water}$ [d] |
|---|---|---|---|---|
| **$\omega$B97X-D3/def2-TZVP** | **-1.3** | – | **15.98** | – |
| PM7//$\omega$B97X-D3/def2-TZVP | -1.82 | – | 23.57 | – |
| PM6//$\omega$B97X-D3/def2-TZVP | 7.1 | – | 29.99 | – |
| PM6-D3//$\omega$B97X-D3/def2-TZVP | 4.94 | – | 27.99 | – |
| PM3//$\omega$B97X-D3/def2-TZVP | -1.26 | – | 29.06 | – |
| **B3LYP-D3BJ/def2-TZVP** | **-2.51** | **3.75** | – | – |
| PM7//B3LYP-D3BJ/def2-TZVP | -3.01 | 2.75 | – | – |
| PM6//B3LYP-D3BJ/def2-TZVP | 7.0 | 12.72 | – | – |
| PM6-D3//B3LYP-D3BJ/def2-TZVP | 4.76 | 10.58 | – | – |
| PM3//B3LYP-D3BJ/def2-TZVP | -1.54 | 4.15 | – | – |
| **BLYP-D3BJ/def2-SVP** | **-1.41** | **3.48** | **17.72** | **20.54** |
| PM7//BLYP-D3BJ/def2-SVP | -2.53 | 3.29 | 20.66 | 22.27 |
| PM6//BLYP-D3BJ/def2-SVP | 5.79 | 11.94 | 27.29 | 28.18 |
| PM6-D3//BLYP-D3BJ/def2-SVP | 3.39 | 9.59 | 25.16 | 26.42 |
| PM3//BLYP-D3BJ/def2-SVP | -1.58 | 4.06 | 28.85 | 28.28 |
| PM7(opt)//BLYP-D3BJ/def2-SVP | -0.85 | 5.27 | 22.4 | 23.57 |
| PM7(opt)//B3LYP-D3BJ/def2-TZVP | -1.37 | 4.31 | – | – |

[a] Enthalpy of the WC geometry formation in gas phase; [b] Enthalpy of the WC geometry formation in implicit water model; [c] Activation enthalpy in gas phase; [d] Activation enthalpy in implicit water model; All enthalpies are given in kcal/mol. The reference DFT entries are highlighted with bold font.

**Supplementary Table S3:** Values of the decoding rate constants used for kinetic modeling

| Designation [a] | value | unit | condition [b] | reference |
|---|---|---|---|---|
| $k_1$ | 140 | $\mu M^{-1}$ $s^{-1}$ | 20 °C, HiFi | Rudorf et al.[S48] |
| $q_2$ | 85 | $s^{-1}$ | 20 °C, HiFi | Rudorf et al.[S48] |
| $k_2$ | 720 [c] | $s^{-1}$ | arbitrary [c] | Pavlov and Ehrenberg[S50] |
| $q_3^c$ | 25 [c] | $s^{-1}$ | arbitrary [c] | Pavlov and Ehrenberg[S50] |
| $q_3^{nc}$ | 3900 [c] | $s^{-1}$ | arbitrary [c] | Pavlov and Ehrenberg[S50] |
| $k_3$ | 180 | $s^{-1}$ | 20 °C, HiFi | Rudorf et al.[S48] |
| $q_4^c$ | 0.2 | $s^{-1}$ | 20 °C, HiFi | Rudorf et al.[S48] |
| $q_4^{nc}$ | 140 | $s^{-1}$ | 20 °C, HiFi | Rudorf et al.[S48] |
| $k_4^c$ | 190 | $s^{-1}$ | 20 °C, HiFi | Rudorf et al.[S48] |
| $k_4^{nc}$ | 0.6 | $s^{-1}$ | 20 °C, HiFi | Rudorf et al.[S48] |

[a] Designations for the rate constants used in our study, as well as in Pavlov and Ehrenberg[S50]. Designations in Rudorf et al.[S48] can be different; [b] Experimental conditions at which the rate constants were measured. HiFi – high-fidelity conditions (3.5 mM $Mg^{2+}$, 0.5 mM spermidine, and 8 mM putrescine)[S76]; [c] The values were obtained from the free energy diagram in Pavlov and Ehrenberg[S50], and uniformly rescaled to obtain values eligible for numerical calculations using ODE.

### Supplementary Figures

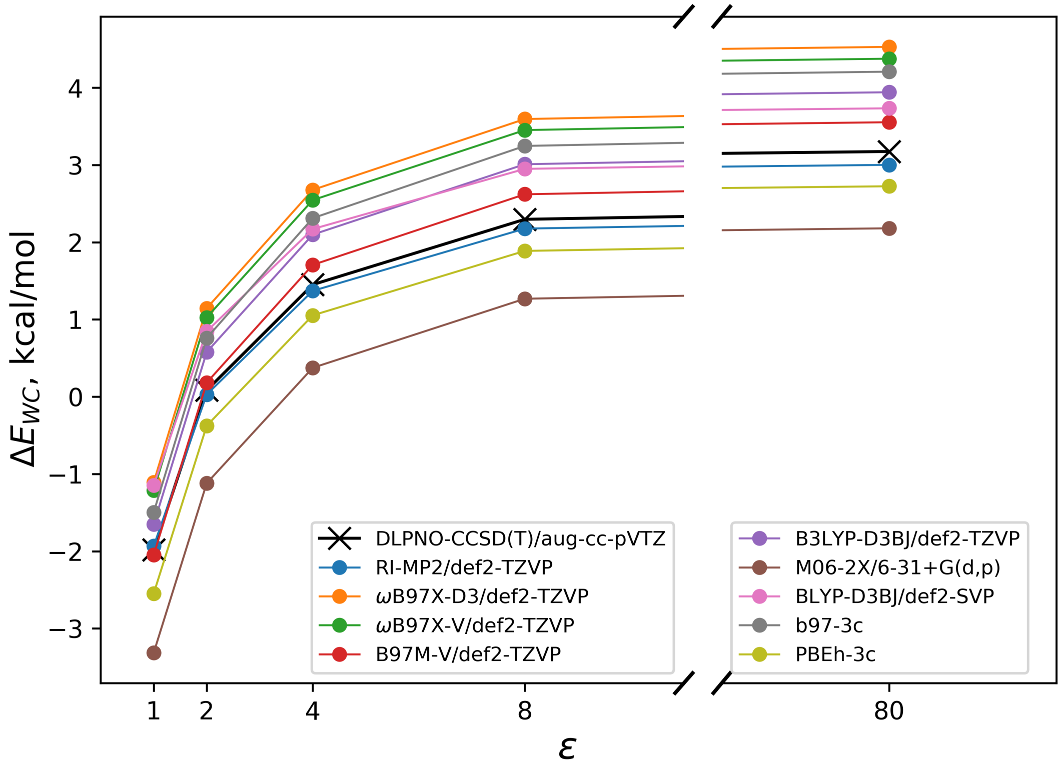


**Supplementary Figure S1:** Dependence of total energy of the WC geometry formation $\Delta E_{wc}$ on dielectric constant $\varepsilon$ of the implicit solvent model.

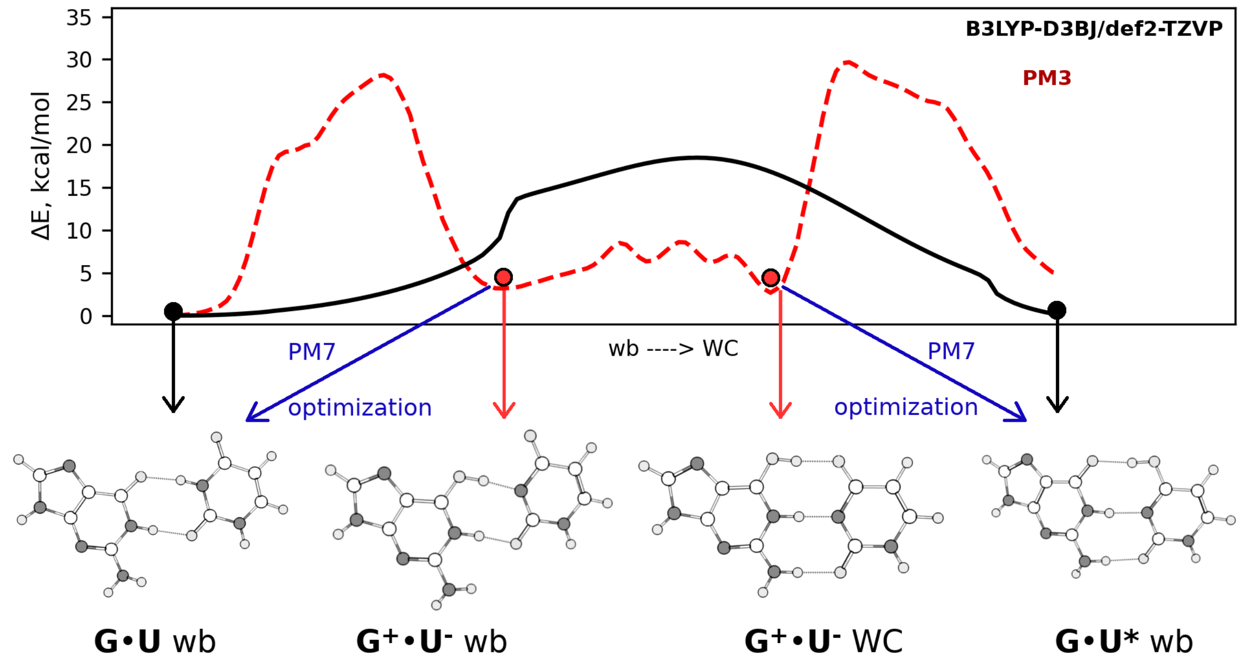


**Supplementary Figure S2:** Spurious wb-WC pathway optimized with PM3 method. The plot shows energy profiles from nudged elastic band optimizations of the wb-WC reaction at DFT (B3LYP-D3BJ/def2-TZVP, black) and US calculations at PM3 (red) levels of theory. The dots on the curves denote local minima. PM3 pathway contains two additional local minima: ion pairs $G^{+}\circ U^{-}$ wb and $G^{+}\circ\ U^{-}$ WC (shown below). Geometry optimizations of these ion pair structures demonstrate that they are not stationary points on the PM7 level (as well as in all tested DFT methods), and converge to G∘U wb and G∘U* WC, respectively.

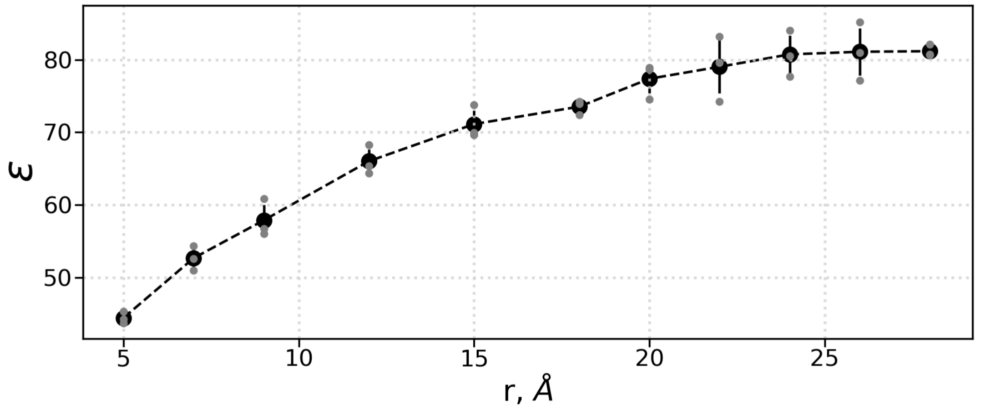


**Supplementary Figure S3:** Size-dependence of dielectric constant $\varepsilon$ calculations in our KFF-based approach applied to the box of 42,000 TIP3P water molecules. Black curve shows the mean values from 3 replicas, while gray circles denote individual replicas.

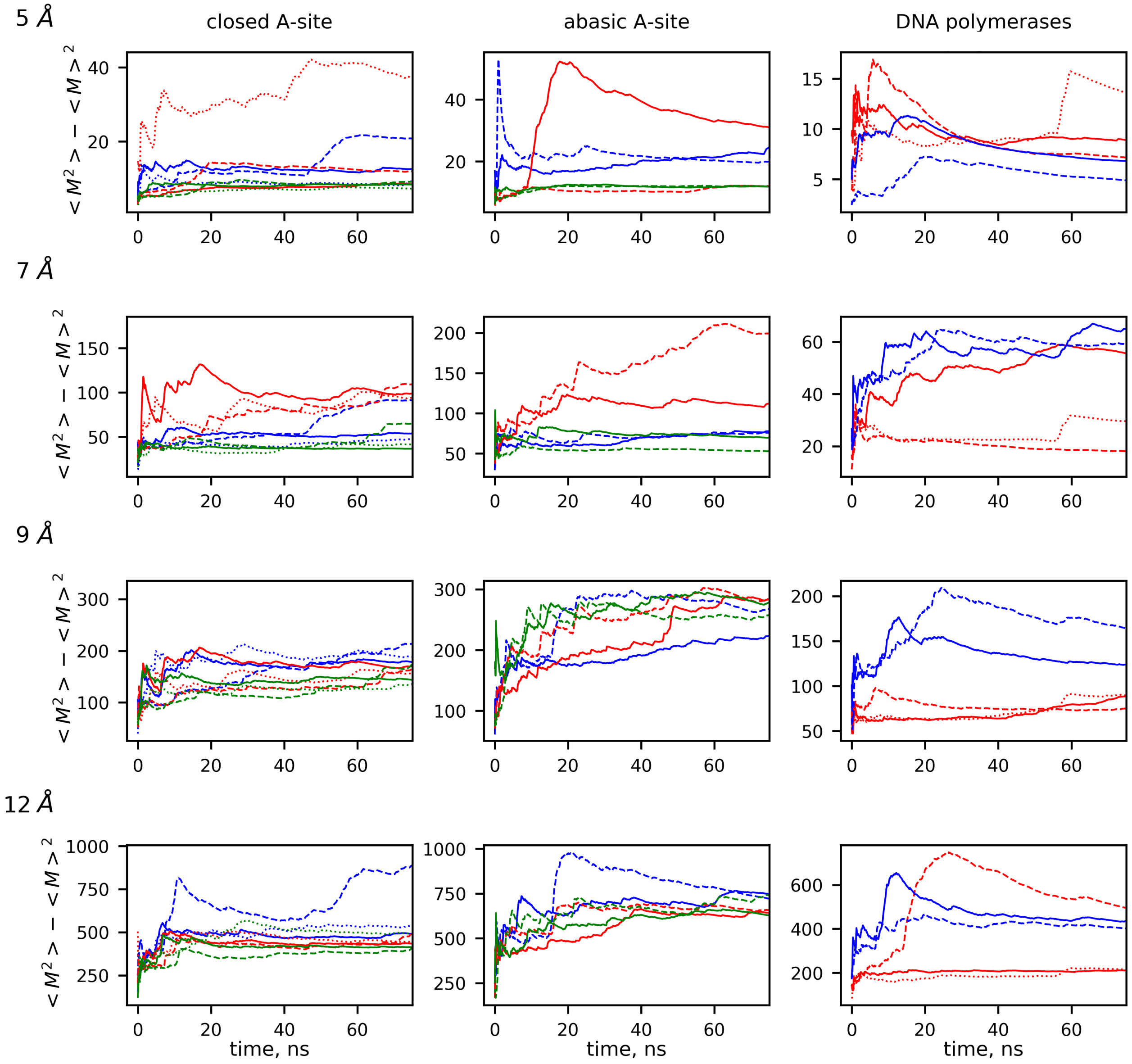


**Supplementary Figure S4:** Cumulative mean square dipole moment fluctuation in all analyzed trajectories. Each row represents a distance cutoff, while each column represents a studied system. For the A-site models, fluctuations of the dipole moment around three codon-anticodon positions are shown on each plot as blue, red and green curves denoting the first, second and third codon-anticodon position, respectively. Solid, dashed and dotted lines denote different replicas. For the DNA polymerases, red and blue curves denote pol-$\beta$ and T7-pol, respectively.

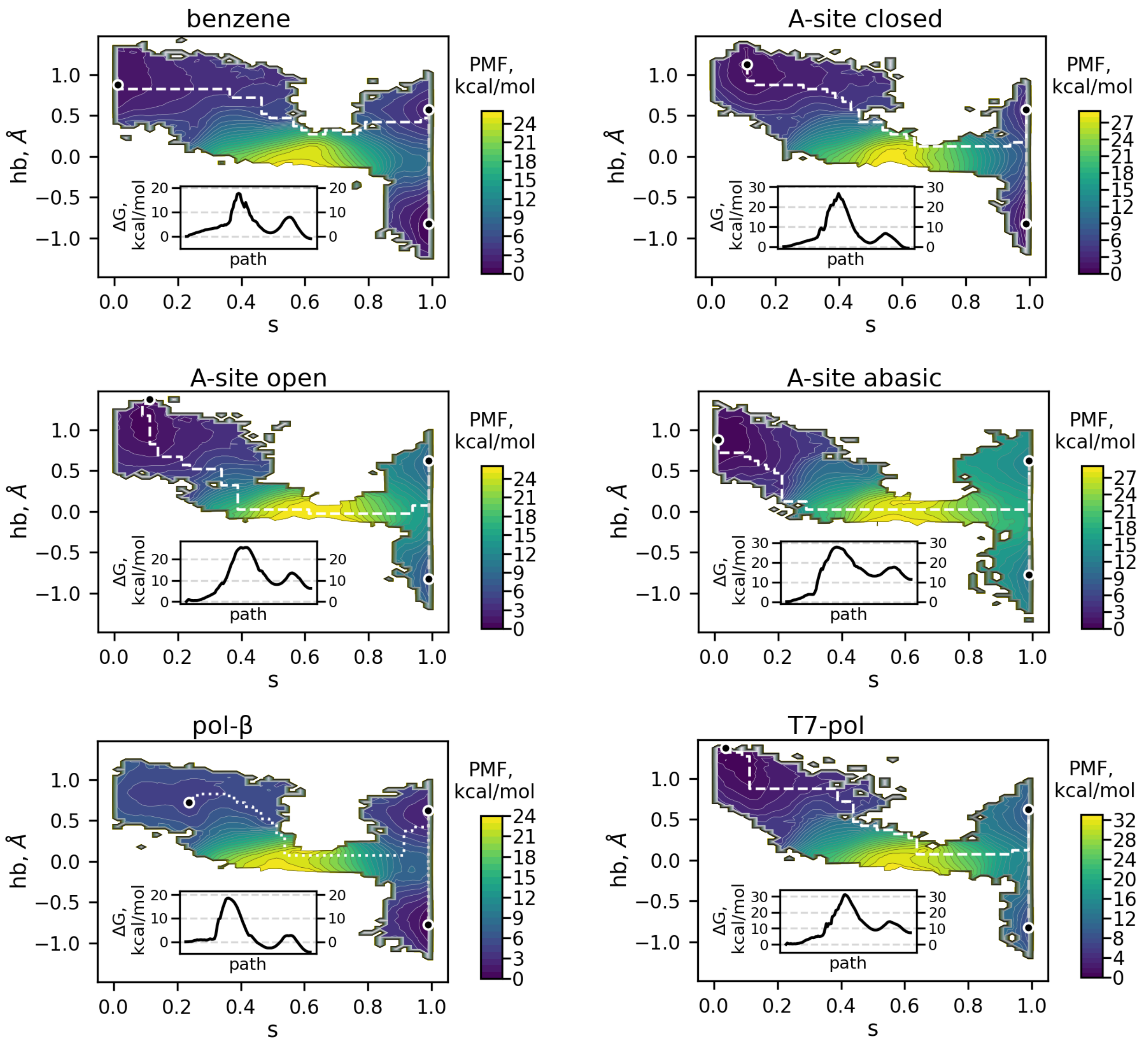


**Supplementary Figure S5:** Final US-derived PMFs of the wb-WC reaction in all systems except DNA heptamer, which is shown on Fig. 4A. See the legend on Fig. 4A.

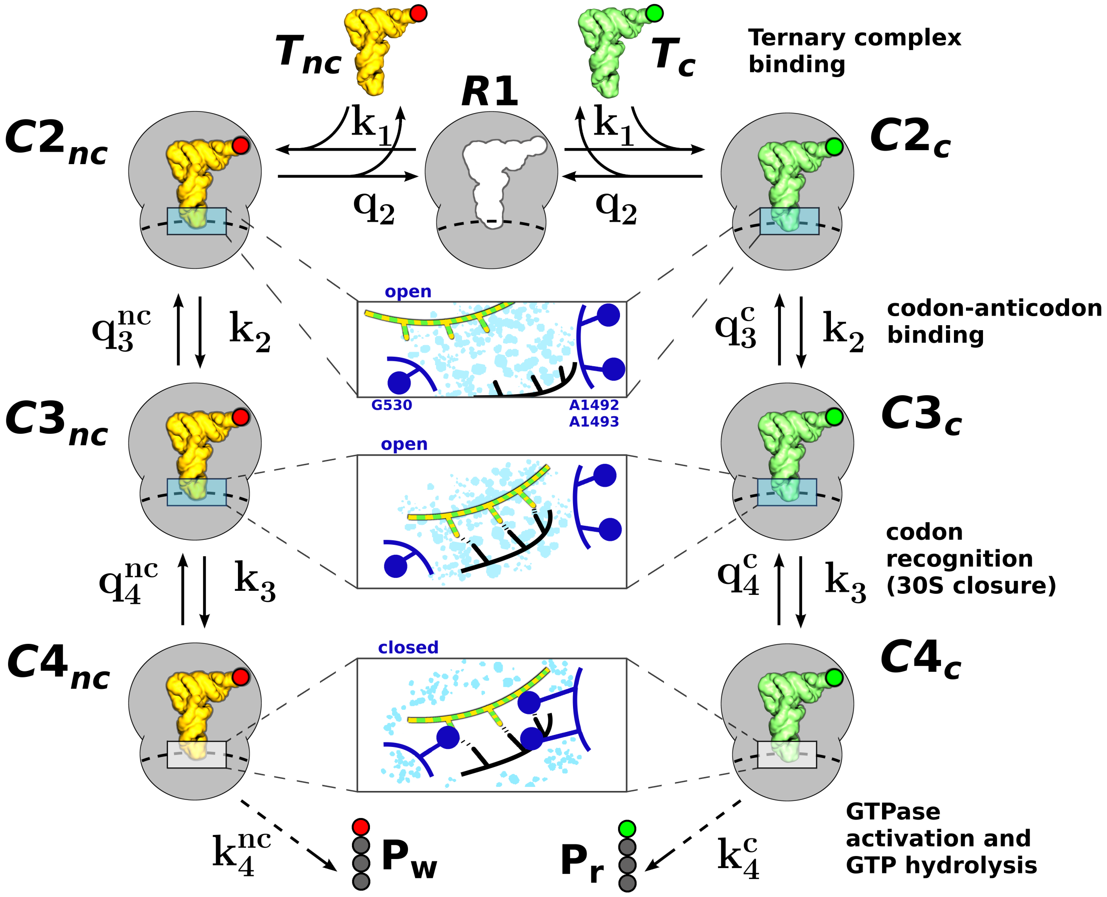


**Supplementary Figure S6:** "Classical" model of initial selection in translation. The nomenclature of the states is described in details in Pavlov and Ehrenberg[S50] and is supported by cryo-EM studies[S60,S77] and kinetic studies[S49]. The steps following the initial selection are not considered in this model, therefore the GTPase activation step leads directly to the peptides. $R1$ represents mRNA-programmed ribosomes with empty A-site. $T_c$ and $T_{nc}$ represent cognate and near-cognate ternary complexes, respectively. In $C2$ state the codon-anticodon helix is not yet formed, and the decoding site is open. In $C3$ the codon-anticodon helix forms in the open decoding site. In $C4$ the decoding site is in the closed state.

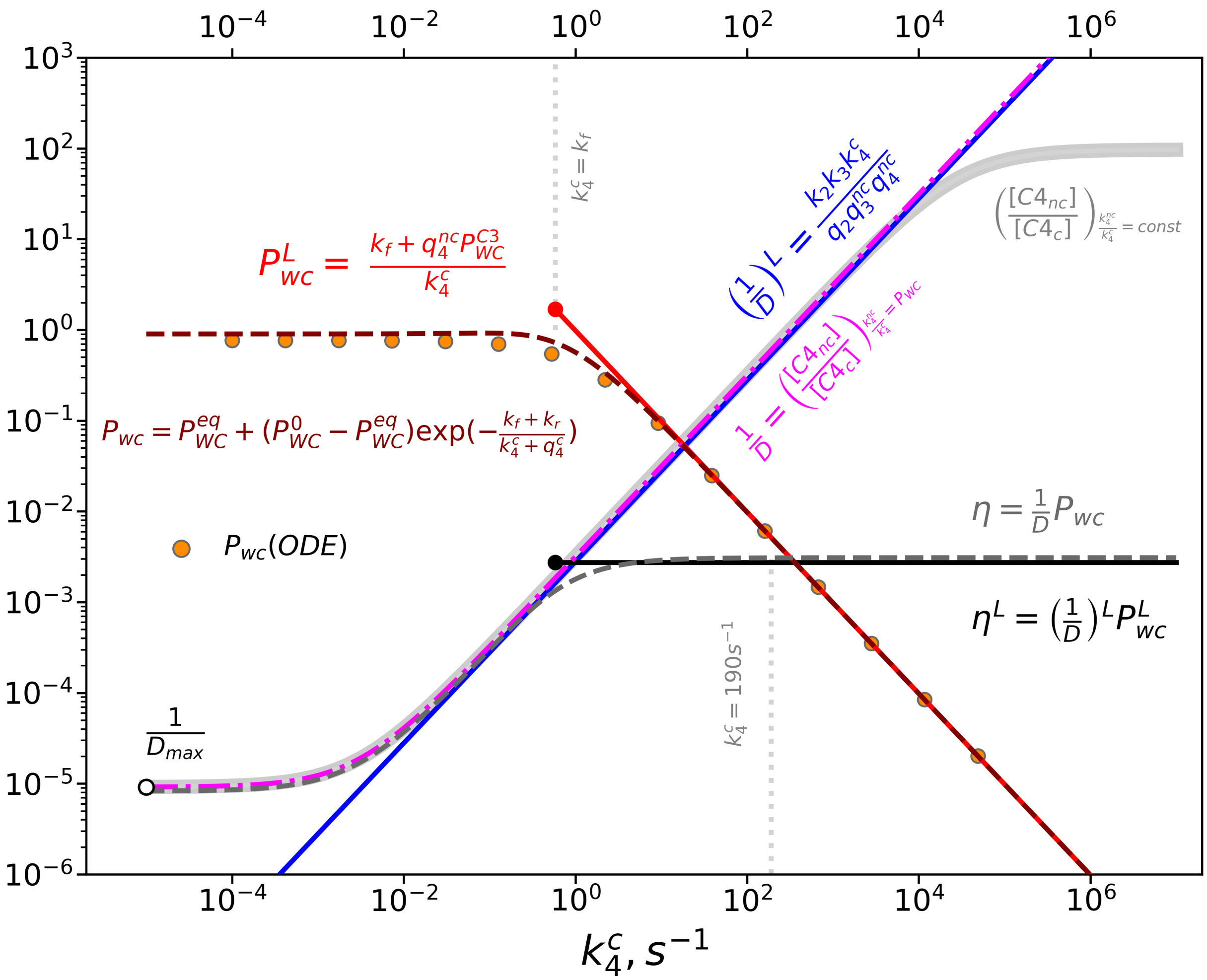


**Supplementary Figure S7:** Visualization of the terms contributing to the flat $\eta(k_4^c)$ curve in the slow kinetic regime of the wb-WC reaction. See Appendix for the derivations of terms shown on the figure. Equations on the plot label curves with the corresponding colors. The orange dots denote the numerical calculations $P_{wc}^{ODE}$. $\left(\frac{1}{D}\right)^L$, $P_{wc}^L$ and $\eta^L$ represent the linear approximations of the corresponding equations. The linear approximations $P_{wc}^L$ and $\eta^L$ are valid only for $k_4^c > k_f$. The wb-WC parameters used for analytical and numerical calculations were ($\Delta G_{wc}^{C4} = -1$ kcal/mol, $\Delta G_{wc}^{C3} = 3.4$, $\Delta G_{\ddagger} = 17.8$ kcal/mol)

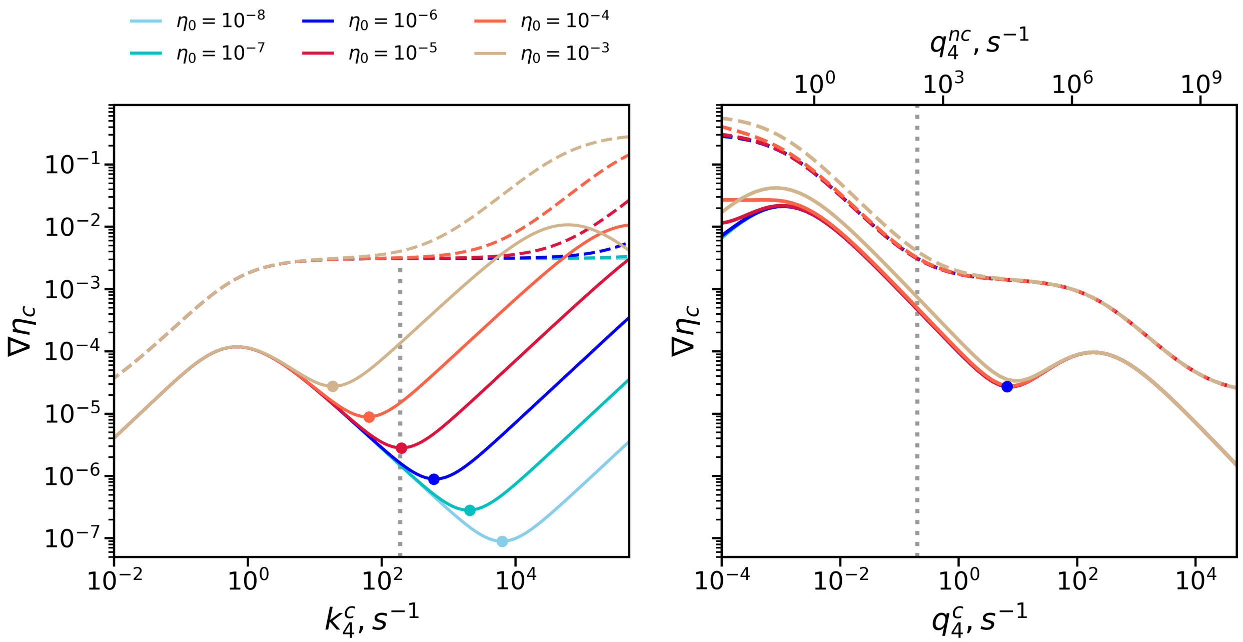


**Supplementary Figure S8:** Cumulative error rate $\eta_c$ and its gradient $\nabla\eta_c$ for a range of scaled down $\eta_0$ curves as functions of $k_4^c$ and $q_4^{c/nc}$. The colors denote the $\eta_0$ values at $k_4^c = 190s^{-1}$ and $q_4^{nc} = 140s^{-1}$ as contributions to $\eta_c$. The dashed and solid lines denote $\eta_c$ and $\nabla\eta_c$, respectively. Local minima in $\nabla\eta_c$ are shown as circles of corresponding colors. The gray vertical lines denote the experimental values of $k_4^c$ and $q_4^{c/nc}$.

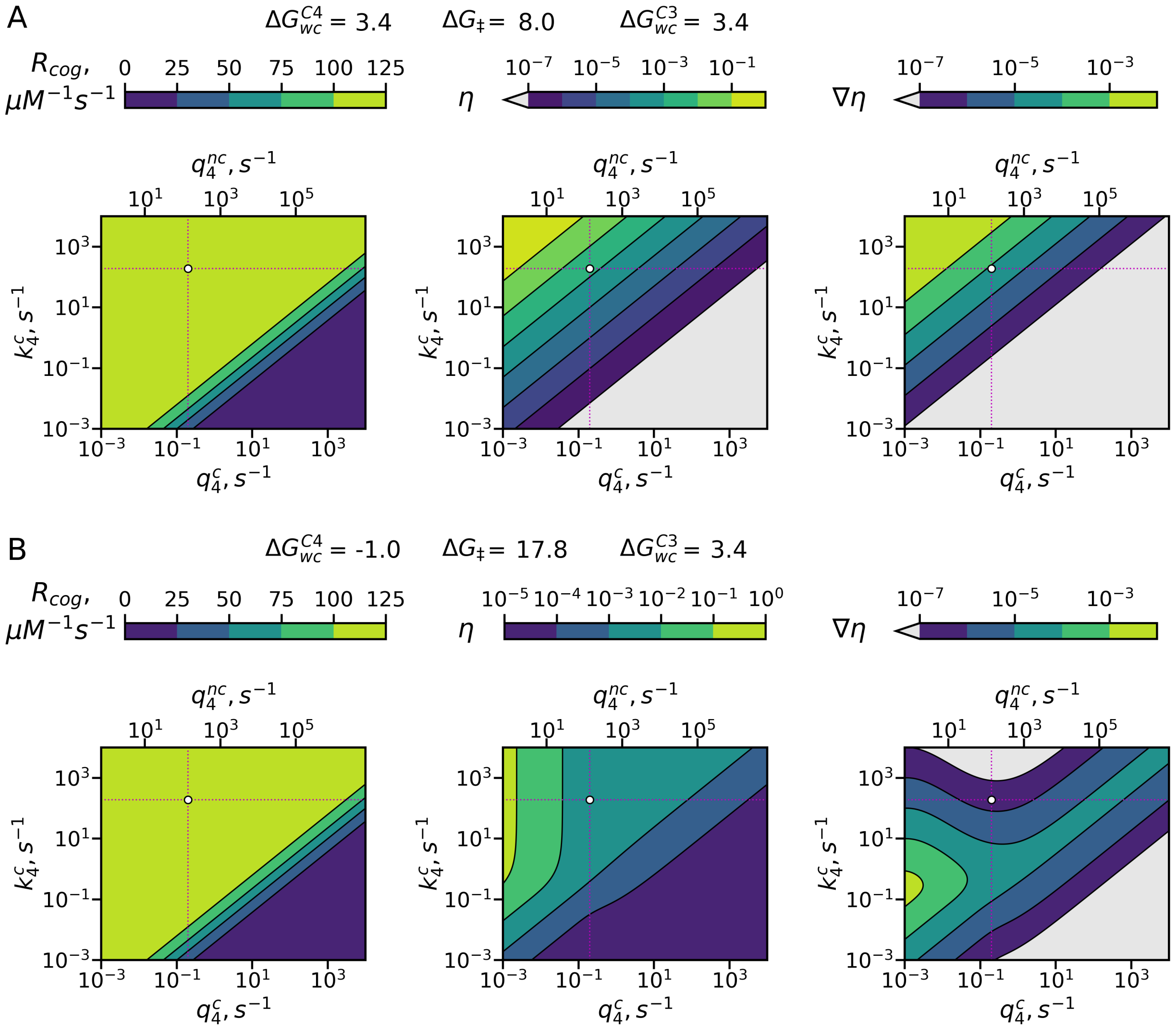


**Supplementary Figure S9:** Optimization landscapes of cognate decoding rate $R_{cog}$, $\eta$ and $\nabla\eta$ for the fast (**A**) and slow (**B**) kinetic regimes of the wb-WC reaction. The wb-WC parameters of the visualized kinetic regimes are shown at the top of the plots. See the captions in Fig. 6 for further information.

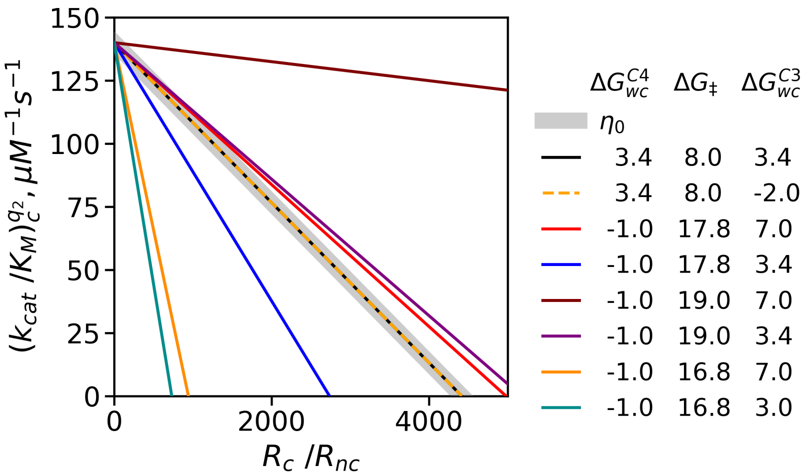


**Supplementary Figure S10:** Linear rate-accuracy trade-offs obtained by varying $q_2$ (modeling the effects of $Mg^{2+}$ concentration). Instead of visualizing $\eta(q_2)$, the plot is designed to be directly comparable to the plots in Zhang et al.[S68]. As can be seen, the linear trade-offs at $q_2$ do not necessarily imply fast tautomeric equilibration throughout all initial selection process, contrary to what suggested by Pavlov et al.[S78]. Instead, kinetic and thermodynamic properties of the wb-WC reaction in C3 and C4 states affect the slope of the straight trade-off lines, similarly to the effects of type and position of a mismatch observed in Zhang et al.[S68]. The visualized wb-WC parameter combinations are shown on the right.

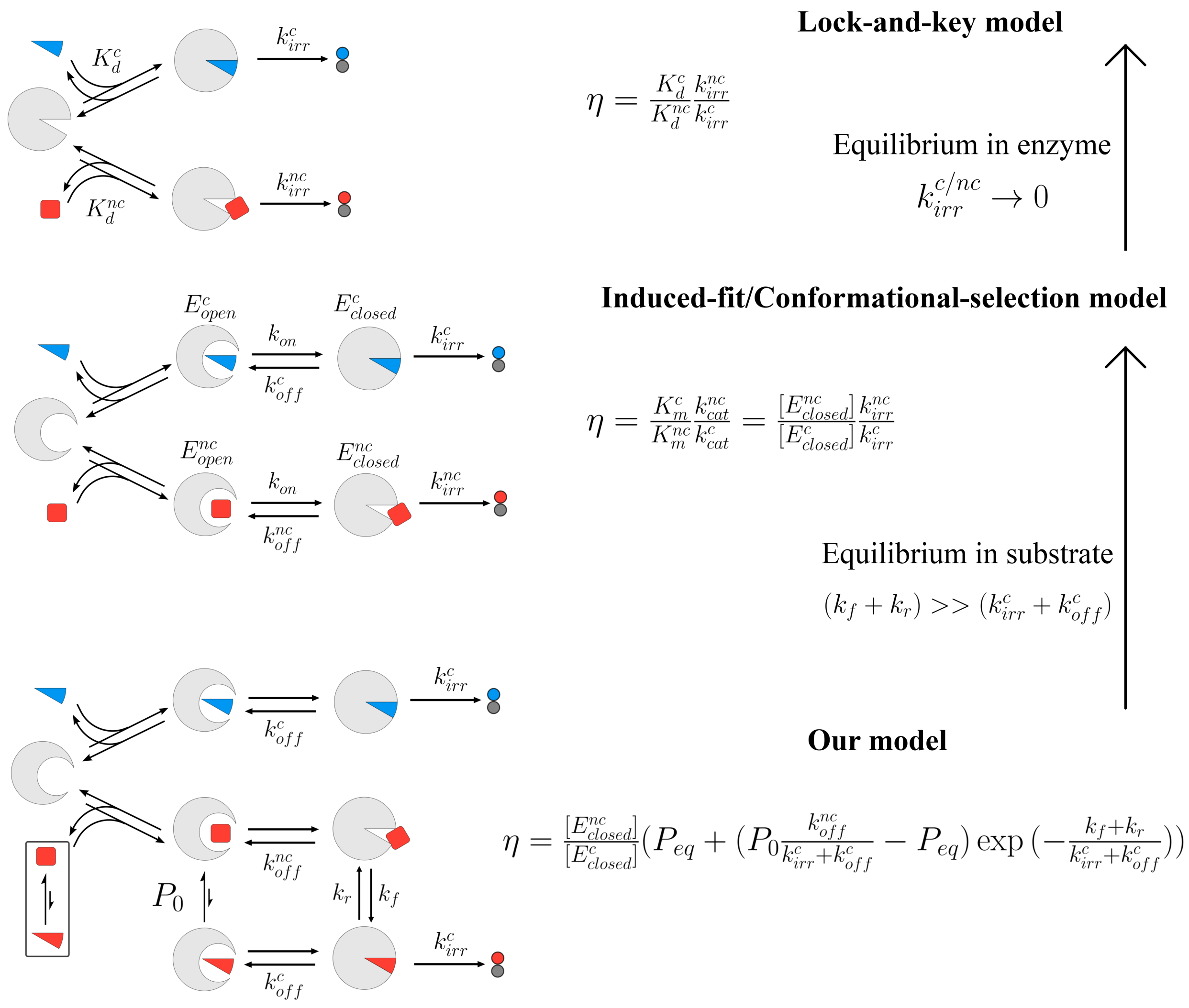


**Supplementary Figure S11:** Simplified schemes of the substrate recognition models.

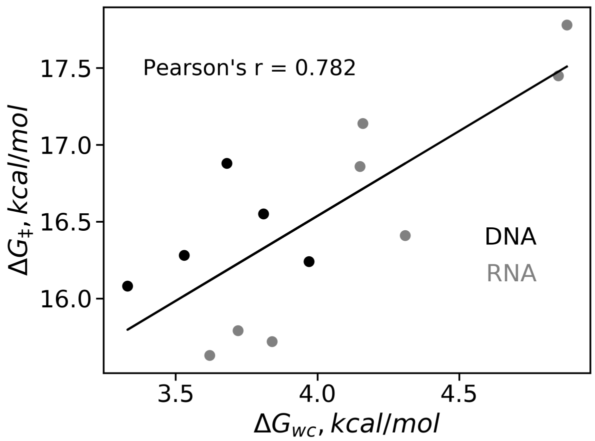


**Supplementary Figure S12:** Linear correlation between $\Delta G_{WC}$ and $\Delta G_{\ddagger}$ of the wb-WC reaction in G∘U(T) of RNA and DNA duplexes in solution, obtained with NMR by Kimsey et al.[S70]. Only neutral pH conditions and unmodified nucleobases were considered. Measurements in DNA and RNA duplexes are shown in black and grey, respectively. The black line denotes the linear fit of all measurements. Pearson's correlation coefficient is shown on the plot.

(S7) Stewart, J. J. Mopac2016. 2016.

(S8) Jorgensen, W. L.; Chandrasekhar, J.; Madura, J. D.; Impey, R. W.; Klein, M. L. Comparison of simple potential functions for simulating liquid water. *The Journal of Chemical Physics* **1983**, *79*, 926–935.

(S9) Koag, M. C.; Nam, K.; Lee, S. The spontaneous replication error and the mismatch discrimination mechanisms of human DNA polymerase $\beta$. *Nucleic Acids Research* **2014**, *42*, 11233–11245.

(S10) Freudenthal, B. D.; Beard, W. A.; Shock, D. D.; Wilson, S. H. Observing a DNA polymerase choose right from wrong. *Cell* **2013**, *154*, 157.

(S11) Doublié, S.; Tabor, S.; Long, A. M.; Richardson, C. C.; Ellenberger, T. Crystal structure of a bacteriophage T7 DNA replication complex at 2.2 Å resolution. *Nature* **1998**, *391*, 251–258.

(S12) Humphrey, W.; Dalke, A.; Schulten, K. VMD - Visual Molecular Dynamics. *Journal of Molecular Graphics* **1996**, *14*, 33–38.

(S13) Rozov, A.; Wolff, P.; Grosjean, H.; Yusupov, M.; Yusupova, G.; Westhof, E. Tautomeric GU pairs within the molecular ribosomal grip and fidelity of decoding in bacteria. *Nucleic Acids Research* **2018**,

(S14) Zeng, X.; Chugh, J.; Casiano-Negroni, A.; Al-Hashimi, H. M.; Brooks Iii, C. L. Flipping of the Ribosomal A-Site Adenines Provides a Basis for tRNA Selection. *Journal of Molecular Biology* **2014**, *426*, 3201–3213.

(S15) Casalino, L.; Palermo, G.; Abdurakhmonova, N.; Rothlisberger, U.; Magistrato, A. Development of site-specific $Mg^{2+}$-RNA force field parameters: A dream or reality? Guidelines from combined molecular dynamics and quantum mechanics simulations. *Journal of Chemical Theory and Computation* **2017**, *13*, 340–352.

(S16) Šponer, J.; Bussi, G.; Krepl, M.; Banáš, P.; Bottaro, S.; Cunha, R. A.; Gil-Ley, A.; Pinamonti, G.; Poblete, S.; Jurečka, P.; Walter, N. G.; Otyepka, M. RNA Structural Dynamics As Captured by Molecular Simulations: A Comprehensive Overview. *Chemical Reviews* **2018**,

(S17) Zheng, H.; Shabalin, I. G.; Handing, K. B.; Bujnicki, J. M.; Minor, W. Magnesium-binding architectures in RNA crystal structures: Validation, binding preferences, classification and motif detection. *Nucleic Acids Research* **2015**, *43*, 3789–3801.

(S18) Nierhaus, K. H. $Mg^{2+}$, $K^+$, and the ribosome. *Journal of Bacteriology* **2014**, *196*, 3817–3819.

(S19) Robbins, T. J.; Wang, Y. Effect of initial ion positions on the interactions of monovalent and divalent ions with a DNA duplex as revealed with atomistic molecular dynamics simulations. *Journal of Biomolecular Structure and Dynamics* **2013**, *31*, 1311–1323.

(S20) Beard, W. A.; Wilson, S. H. Structure and mechanism of DNA polymerase $\beta$. *Chemical Reviews* **2006**, *106*, 361–382.

(S21) Ogle, J. M.; Murphy IV, F. V.; Tarry, M. J.; Ramakrishnan, V. Selection of tRNA by the ribosome requires a transition from an open to a closed form. *Cell* **2002**, *111*, 721–732.

(S22) Xu, Y.; Vanommeslaeghe, K.; Aleksandrov, A.; MacKerell, A. D.; Nilsson, L. Additive CHARMM force field for naturally occurring modified ribonucleotides. *Journal of Computational Chemistry* **2016**, *37*, 896–912.

(S23) Phillips, J. C.; Hardy, D. J.; Maia, J. D.; Stone, J. E.; Ribeiro, J. V.; Bernardi, R. C.; Buch, R.; Fiorin, G.; Hénin, J.; Jiang, W.; McGreevy, R.; Melo, M. C.; Radak, B. K.; Skeel, R. D.; Singharoy, A.;

Wang, Y.; Roux, B.; Aksimentiev, A.; Luthey-Schulten, Z.; Kalé, L. V.; Schulten, K.; Chipot, C.; Tajkhorshid, E. Scalable molecular dynamics on CPU and GPU architectures with NAMD. *Journal of Chemical Physics* **2020**, *153*, 044130.

(S24) Huang, J.; Mackerell, A. D. CHARMM36 all-atom additive protein force field: Validation based on comparison to NMR data. *Journal of Computational Chemistry* **2013**, *34*, 2135–2145.

(S25) Best, R. B.; Zhu, X.; Shim, J.; Lopes, P. E. M.; Mittal, J.; Feig, M.; MacKerell, A. D. Optimization of the Additive CHARMM All-Atom Protein Force Field Targeting Improved Sampling of the Backbone $\phi$, $\psi$ and Side-Chain $\chi1$ and $\chi2$ Dihedral Angles. *Journal of Chemical Theory and Computation* **2012**, *8*, 3257–3273.

(S26) Denning, E. J.; Priyakumar, U. D.; Nilsson, L.; Mackerell, A. D. Impact of 2′-hydroxyl sampling on the conformational properties of RNA: Update of the CHARMM all-atom additive force field for RNA. *Journal of Computational Chemistry* **2011**, *32*, 1929–1943.

(S27) Darden, T.; York, D.; Pedersen, L. Particle mesh Ewald: An N·log(N) method for Ewald sums in large systems. *The Journal of Chemical Physics* **1993**, *98*, 10089–10092.

(S28) Miyamoto, S.; Kollman, P. A. Settle: An analytical version of the SHAKE and RATTLE algorithm for rigid water models. *Journal of Computational Chemistry* **1992**, *13*, 952–962.

(S29) Andersen, H. C. Rattle: A ”velocity” version of the shake algorithm for molecular dynamics calculations. *Journal of Computational Physics* **1983**, *52*, 24–34.

(S30) Pitera, J. W.; Falta, M.; Van Gunsteren, W. F. Dielectric properties of proteins from simulation: The effects of solvent, ligands, pH, and temperature. *Biophysical Journal* **2001**, *80*, 2546–2555.

(S31) Kolafa, J.; Viererblová, L. Static dielectric constant from simulations revisited: Fluctuations or external field? *Journal of Chemical Theory and Computation* **2014**, *10*, 1468–1476.

(S32) Yang, L.; Weerasinghe, S.; Smith, P. E.; Pettitt, B. M. Dielectric response of triplex DNA in ionic solution from simulations. *Biophysical Journal* **1995**, *69*, 1519–1527.

(S33) Chandra, A. Static dielectric constant of aqueous electrolyte solutions: is there any dynamic contribution? *Journal of Chemical Physics* **2000**, *113*, 903–905.

(S34) Gereben, O.; Pusztai, L. On the accurate calculation of the dielectric constant from molecular dynamics simulations: The case of SPC/E and SWM4-DP water. *Chemical Physics Letters* **2011**, *507*, 80–83.

(S35) Branduardi, D.; Gervasio, F. L.; Parrinello, M. From A to B in free energy space. *Journal of Chemical Physics* **2007**, *126*, 054103.

(S36) Fiorin, G.; Klein, M. L.; Hénin, J. Using collective variables to drive molecular dynamics simulations. *Molecular Physics* **2013**, *111*, 3345–3362.

(S37) Henkelman, G.; Uberuaga, B. P.; Jónsson, H. Climbing image nudged elastic band method for finding saddle points and minimum energy paths. *Journal of Chemical Physics* **2000**, *113*, 9901–9904.

(S38) Brovarets, O. O.; Hovorun, D. M. How many tautomerization pathways connect Watson-Crick-like G*·T DNA base mispair and wobble mismatches? *J Biomol Struct Dyn* **2015**, 1–19.

(S39) Kästner, J. Umbrella sampling. *Wiley Interdisciplinary Reviews: Computational Molecular Science* **2011**, *1*, 932–942.

(S40) Melo, M. C. R.; Bernardi, R. C.; Rudack, T.; Scheurer, M.; Riplinger, C.; Phillips, J. C.; Maia, J. D. C.; Rocha, G. B.; Ribeiro, J. V.; Stone, J. E.; Neese, F.; Schulten, K.; Luthey-Schulten, Z. NAMD goes quantum: an integrative suite for hybrid simulations. *Nature Methods* **2018**, *15*, 351–354.

(S41) Marcos-Alcalde, I.; Setoain, J.; Mendieta-Moreno, J. I.; Mendieta, J.; Gómez-Puertas, P. MEPSA: Minimum energy pathway analysis for energy landscapes. *Bioinformatics* **2015**, *31*, 3853–3855.

(S42) Pape, T.; Wintermeyer, W.; Rodnina, M. V. Conformational switch in the decoding region of 16S rRNA during aminoacyl-tRNA selection on the ribosome. *Nature Structural Biology* **2000**, *7*, 104–107.

(S43) Pernod, K.; Schaeffer, L.; Chicher, J.; Hok, E.; Rick, C.; Geslain, R.; Eriani, G.; Westhof, E.; Ryckelynck, M.; Martin, F. The nature of the purine at position 34 in tRNAs of 4-codon boxes is correlated with nucleotides at positions 32 and 38 to maintain decoding fidelity. *Nucleic acids research* **2020**, *48*, 6170–6183.

(S44) Garofalo, R.; Wohlgemuth, I.; Pearson, M.; Lenz, C.; Urlaub, H.; Rodnina, M. V. Broad range of missense error frequencies in cellular proteins. *Nucleic acids research* **2019**, *47*, 2932–2945.

(S45) Mordret, E.; Dahan, O.; Asraf, O.; Rak, R.; Yehonadav, A.; Barnabas, G. D.; Cox, J.; Geiger, T.; Lindner, A. B.; Pilpel, Y. Systematic Detection of Amino Acid Substitutions in Proteomes Reveals Mechanistic Basis of Ribosome Errors and Selection for Translation Fidelity. *Molecular Cell* **2019**,

(S46) Manickam, N.; Nag, N.; Abbasi, A.; Patel, K.; Farabaugh, P. J. Studies of translational misreading in vivo show that the ribosome very efficiently discriminates against most potential errors. *RNA* **2014**, *20*, 9–15.

(S47) Zhang, Z.; Shah, B.; Bondarenko, P. V. G/U and Certain Wobble Position Mismatches as Possible Main Causes of Amino Acid Misincorporations. *Biochemistry* **2013**, *52*, 8165–8176.

(S48) Rudorf, S.; Thommen, M.; Rodnina, M. V.; Lipowsky, R. Deducing the kinetics of protein synthesis in vivo from the transition rates measured in vitro. *PLoS Comput Biol* **2014**, *10*, e1003909.

(S49) Rodnina, M. V.; Fischer, N.; Maracci, C.; Stark, H. Ribosome dynamics during decoding. *Philosophical Transactions of the Royal Society B: Biological Sciences* **2017**, *372*.

(S50) Pavlov, M. Y.; Ehrenberg, M. Substrate-Induced Formation of Ribosomal Decoding Center for Accurate and Rapid Genetic Code Translation. *Annual Review of Biophysics* **2018**, *47*, 525–548.

(S51) Harris, C. R.; Millman, K. J.; van der Walt, S. J.; Gommers, R.; Virtanen, P.; Cournapeau, D.; Wieser, E.; Taylor, J.; Berg, S.; Smith, N. J.; Kern, R.; Picus, M.; Hoyer, S.; van Kerkwijk, M. H.; Brett, M.; Haldane, A.; del Río, J. F.; Wiebe, M.; Peterson, P.; Gérard-Marchant, P.; Sheppard, K.; Reddy, T.; Weckesser, W.; Abbasi, H.; Gohlke, C.; Oliphant, T. E. Array Programming with NumPy. *Nature* **2020**, *585*, 357.

(S52) Hunter, J. D. Matplotlib: A 2D graphics environment. *Computing in Science and Engineering* **2007**, *9*, 90–95.

(S53) Fischer, E. Einfluss der Configuration auf die Wirkung der Enzyme. *Berichte der deutschen chemischen Gesellschaft* **1894**, *27*, 2985–2993.

(S54) Koshland, D. E. Enzyme flexibility and enzyme action. *Journal of cellular and comparative physiology* **1959**, *54*, 245–258.

(S55) Changeux, J. P.; Edelstein, S. Conformational selection or induced fit? 50 Years of debate resolved. 2011; http://www.f1000.com/reports/b/3/19.

(S56) Bebenek, K.; Pedersen, L. C.; Kunkel, T. A. Replication infidelity via a mismatch with WatsonCrick geometry. *Proceedings of the National Academy of Sciences* **2011**, *108*, 1862–1867.

(S57) Demeshkina, N.; Jenner, L.; Westhof, E.; Yusupov, M.; Yusupova, G. A new understanding of the decoding principle on the ribosome. *Nature* **2012**, *484*, 256–259.

(S58) Demeshkina, N.; Jenner, L.; Westhof, E.; Yusupov, M.; Yusupova, G. New structural insights into the decoding mechanism: translation infidelity via a G.U pair with Watson-Crick geometry. *FEBS Lett* **2013**, *587*, 1848–1857.

(S59) Rozov, A.; Demeshkina, N.; Westhof, E.; Yusupov, M.; Yusupova, G. Structural insights into the translational infidelity mechanism. *Nat Commun* **2015**, *6*, 7251.

(S60) Loveland, A. B.; Demo, G.; Grigorieff, N.; Korostelev, A. A. Ensemble cryo-EM elucidates the mechanism of translation fidelity. *Nature* **2017**, *546*, 113–117.

(S61) Johansson, M.; Lovmar, M.; Ehrenberg, M. Rate and accuracy of bacterial protein synthesis revisited. *Current Opinion in Microbiology* **2008**, *11*, 141–147.

(S62) Bohr, N. Über die Serienspektra der Elemente. *Zeitschrift für Physik* **1920**, *2*, 423–469.

(S63) Savir, Y.; Tlusty, T. The Ribosome as an Optimal Decoder: A Lesson in Molecular Recognition. *Cell* **2013**, *153*, 471–479.

(S64) Wohlgemuth, I.; Pohl, C.; Mittelstaet, J.; Konevega, A. L.; Rodnina, M. V. Evolutionary optimization of speed and accuracy of decoding on the ribosome. *Philosophical Transactions of the Royal Society B: Biological Sciences* **2011**, *366*, 2979–2986.

(S65) Thompson, R. C.; Karim, A. M. The accuracy of protein biosynthesis is limited by its speed: High fidelity selection by ribosomes of aminoacyl-tRNA ternary complexes containing GTP[$\gamma$S]. *Proceedings of the National Academy of Sciences of the United States of America* **1982**, *79*, 4922–4926.

(S66) Zeidler, W.; Egle, C.; Ribeiro, S.; Wagner, A.; Katunin, V.; Kreutzer, R.; Rodnina, M.; Wintermeyer, W.; Sprinzl, M. SiteDirected Mutagenesis of Thermus thermophilus Elongation Factor Tu: Replacement of His85, Asp81 and Arg300. *European Journal of Biochemistry* **1995**, *229*, 596–604.

(S67) Zaher, H. S.; Green, R. Hyperaccurate and Error-Prone Ribosomes Exploit Distinct Mechanisms during tRNA Selection. *Molecular Cell* **2010**, *39*, 110–120.

(S68) Zhang, J.; Pavlov, M. Y.; Ehrenberg, M. Accuracy of genetic code translation and its orthogonal corruption by aminoglycosides and $Mg^{2+}$ ions. *Nucleic Acids Research* **2018**, *46*, 1362–1374.

(S69) Hoffer, E. D.; Maehigashi, T.; Fredrick, K.; Dunham, C. M. Ribosomal ambiguity (ram) mutations promote the open (off) to closed (on) transition and thereby increase miscoding. *Nucleic Acids Research* **2019**, *47*, 1557–1563.

(S70) Kimsey, I. J.; Szymanski, E. S.; Zahurancik, W. J.; Shakya, A.; Xue, Y.; Chu, C. C.; Sathyamoorthy, B.; Suo, Z.; Al-Hashimi, H. M. Dynamic basis for dG· dT misincorporation via tautomerization and ionization. *Nature* **2018**, *554*, 195–201.

(S71) Loveland, A. B.; Demo, G.; Korostelev, A. A. Cryo-EM of elongating ribosome with EF-Tu•GTP elucidates tRNA proofreading. *Nature* **2020**, 1–6.

(S72) Ehrenberg, M.; Kurland, C. G.; Ruusala, T. Counting cycles of EF-Tu to measure proofreading in translation. *Biochimie* **1986**, *68*, 261–273.

(S73) Morse, J. C.; Girodat, D.; Burnett, B. J.; Holm, M.; Altman, R. B.; Sanbonmatsu, K. Y.; Wieden, H.-J.; Blanchard, S. C. Elongation factor-Tu can repetitively engage aminoacyl-tRNA within the ribosome during the proofreading stage of tRNA selection. *Proceedings of the National Academy of Sciences* **2020**, 201904469.

(S74) Tsai, Y. C.; Johnson, K. A. A new paradigm for DNA polymerase specificity. *Biochemistry* **2006**, *45*, 9675–9687.

(S75) Sartori, P.; Pigolotti, S. Kinetic versus Energetic Discrimination in Biological Copying. *Physical Review Letters* **2013**, *110*, 188101.

(S76) Gromadski, K. B.; Daviter, T.; Rodnina, M. V. A uniform response to mismatches in codon-anticodon complexes ensures ribosomal fidelity. *Molecular Cell* **2006**, *21*, 369–377.

(S77) Fislage, M.; Zhang, J.; Brown, Z. P.; Mandava, C. S.; Frank, J.; Sanyal, S. Cryo-EM shows stages of initial codon selection on the ribosome by aa-tRNA in ternary complex with GTP and the GTPase-deficient EF-Tu H84A. *Nucleic Acids Research* **2018**, 1–14.

(S78) Pavlov, M. Y.; Liljas, A.; Ehrenberg, M. A recent intermezzo at the Ribosome Club. *Philosophical Transactions of the Royal Society B: Biological Sciences* **2017**, *372*.
